# Programming T cells for Early Cancer Detection with Customized Protease-Activatable Receptors

**DOI:** 10.64898/2026.01.16.699939

**Authors:** Hathaichanok Phuengkham, Ying Chen, Anirudh Sivakumar, Ali H. Zamat, Lena Gamboa, Quoc D. Mac, Hee Jun Lee, Leonard C. Rogers, Jicheng You, Sean A. Steele, Sichen Zhu, Matthew S. Gollins, John Blazeck, Peng Qiu, Gabriel A. Kwong

## Abstract

Early cancer detection has the potential to reduce cancer mortality, yet endogenous tumor-shed biomarkers lack sensitivity for early-stage disease. We report OncoSCOUT, a cancer detection strategy using T cells engineered with protease-activatable receptors (PARs) that conditionally recognize tumor cells and release a synthetic biomarker for detection in urine. These PARs comprise masked synthetic Notch receptors in which antigen binding is blocked by a peptide mimotope tethered via a protease-cleavable linker. We demonstrate that requiring both extracellular protease activity and tumor antigen recognition improves spatial specificity and minimizes off-tumor activation of PAR T cells *in vivo*. To identify tumor-selective PARs, we adoptively transferred a HER2-targeted PAR library displaying ∼160,000 unique 4-mer amino acid linkers and discovered multiple variants significantly enriched in a HER2-positive cancer xenograft model. Using a single customized PAR, we show that OncoSCOUT can detect total tumor burdens as small as 10–30 mm^3^ with significantly improved sensitivity than the protein biomarker CA 15-3 or a 20-plex circulating tumor DNA (ctDNA) assay.

## Introduction

Detecting cancer at an early stage while it is still localized has the potential to reduce cancer-related deaths and improve the likelihood of curative treatment^1^. Yet apart from several established screening methods (e.g., mammograms, colonoscopies, and Papanicolaou tests^2,3^), early detection strategies remain unavailable for most cancer types. Current efforts that primarily rely on analyzing endogenous tumor-derived biomarkers in blood – such as proteins^4,5^, circulating tumor cells^6^, cell-free DNA (cfDNA)^7^, and cancer exosomes^8,9^ – are limited in sensitivity as biomarker levels are scarce at the earliest stages of cancer. For example, multi-cancer tests that detect cfDNA methylation patterns have reported Stage I-II sensitivities ranging from ∼10–30% when deployed in large (>6,000) asymptomatic cohorts^10,11^.

While advances in endogenous biomarker collection^12,13^, detection technologies^14–17^ and analysis^18–20^ are at the forefront and hold promise for improving sensitivity, a parallel strategy involves designing biosensors for systemic delivery that can probe tissues for cancer-associated hallmarks^21^ and, upon detecting them, drive the production of a synthetic biomarker to levels that are readily measurable, enabling earlier detection^22^. Within this framework, cancer detection has been demonstrated using molecular probes that either enhance local imaging contrast in response to dysregulated enzymatic activity^23–25^ or release a synthetic biomarker for detection at a distal site (e.g., a serum^26^, breath^27–29^ or urine^30–32^ sample). Genetically engineered cells have also been reported as living biosensors designed for synthetic biomarker release, including probiotics^33,34^ that exploit immune-privilege within the tumor core and macrophages that undergo tumor-induced phenotypic polarization^35^ as mechanisms for on-tumor activation. While promising, surpassing the detection limits of endogenous biomarkers will require synthetic biomarker strategies with significantly improved molecular specificity and logic-gated mechanisms^36–39^ to minimize off-tumor background while driving on-tumor activation *in vivo*.

Based on the ubiquitous dysregulation of cancer-associated proteases^21,40^, we report an approach for early cancer detection via <u>s</u>ynthetic biomarkers of protease <u>c</u>leavage enabled by <u>o</u>n-target <u>u</u>nmasking of <u>T</u> cells (OncoSCOUT). OncoSCOUT uses T cells engineered with protease-activatable receptors (PARs), which integrate masked single-chain variable fragments (scFv) with synthetic Notch (synNotch) signaling to enhance on-tumor selectivity by requiring both cancer-associated protease activity and antigen recognition for activation. PARs are initially masked by a peptide mimotope linked via a protease-cleavable substrate that following proteolysis, permits antigen binding, synNotch activation and production of a bio-orthogonal synthetic biomarker. We develop an *in vivo* discovery pipeline to perform deep profiling of the tumor substrate repertoire by the adoptive transfer of a PAR T cell library displaying ∼160,000 substrate linkers into mice bearing breast cancer xenograft tumors. Using a PAR incorporating an *in vivo*–selected substrate, we customize OncoSCOUT to detect tumor burdens as small as 10-30 mm³ via a secreted synthetic urinary biomarker, significantly improving sensitivity and specificity compared to the FDA-approved blood biomarker CA 15-3 or a 20-plex ctDNA assay. Our results support that programming T cells with customized PARs can significantly reduce cancer detection thresholds compared to those achievable with endogenous biomarkers.

## Results

### PAR T cell activation requires extracellular protease activity and antigen recognition

We designed PARs by incorporating a peptide-mimotope masked scFv with the previously described design of a synNotch receptor^41,42^ (**Fig. 1A** and **Supplementary Fig. 1A**). We postulated that cleavage of the substrate linker by an extracellular protease would release the peptide mimotope to unmask the scFv for binding, thereby triggering release of the Gal4-VP64 transcription factor to drive reporter expression through the UAS promoter (**Fig. 1B**). To test this, we cloned PARs using scFv sequences derived from anti-human epidermal growth factor receptor 2 (αHER2; trastuzumab^43^) and anti-human epidermal growth factor receptor (αhEGFR; cetuximab^44^), along with their respective validated peptide mimotopes (LLGPYELWELSH^45^ and QGQSGQCISPRGCPDGPYVMY^46^). For each PAR, we generated constructs using either a substrate linker (LVPRGSG^30,47^) selective for the protease thrombin (Thrb) or a flexible GSGGSG linker^46^ as a negative control (**Fig. 1C** and **Supplementary Fig. 2**). In co-culture studies, Thrb-activatable HER2 or hEGFR PAR T cells were only activated when both Thrb and HER2-positive cancer cells were present together, whereas control T cells displaying GSGGSG PARs showed no activation under all tested conditions (**Fig. 1C** and **Supplementary Fig. 2**). Reporter expression depended on the number of HER2-positive tumor cells in co-culture (**Supplementary Fig. 1B**), the concentration of Thrb in the supernatant (**Supplementary Fig. 1C**), and the proteolytic activity of extracellular Thrb, as both the binding of recombinant HER2 to PARs (**Fig. 1D**) and downstream reporter expression (**Fig. 1E**) were disrupted by bivalirudin, a peptide inhibitor of Thrb.

**Figure 1.**
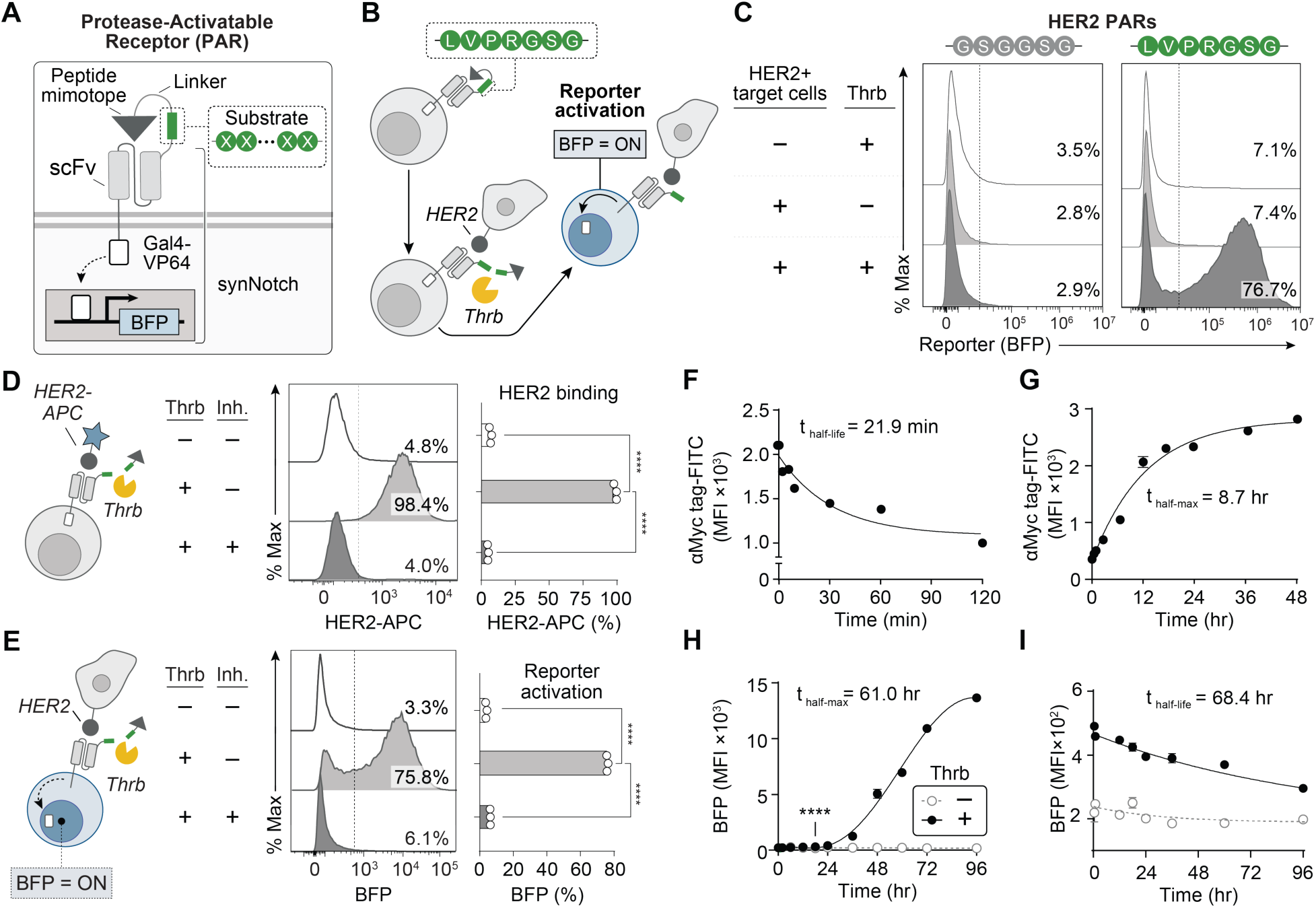
Activation of HER2 PAR T cells requires extracellular protease activity and antigen recognition. (**A**) Schematic of a PAR featuring a masked scFv combined with a synNotch receptor, where antigen binding is blocked by a peptide mimotope linked through a substrate linker. (**B**) After linker proteolysis, scFv engagement triggers release of the transcription factor Gal4-VP64 to drive BFP reporter expression by the UAS promoter. Substrate linker sequence LVPRGSG is cleavable by thrombin (Thrb). (**C**) Flow histogram of BFP expression by primary human T cells engineered with PAR, displaying either a substrate linker cleavable by thrombin (Thrb; LVPRGSG) or a control linker (GSGGSG), after co-incubation with both Thrb and HER2-positive MDA-MB-468 breast cancer cells at a 1:1 cell ratio. (**D**) Representative flow histogram and frequency bar plot of Thrb-activatable PAR T cells stained with recombinant HER2 after co-incubation with Thrb, or Thrb and its inhibitor (inh.) bivalirudin for 30 min at 37°C. (**E**) Representative flow histogram and frequency bar plot of BFP expression by PAR T cells co-incubated with HER2+ MDA-MB-468 cells at a 1:1 cell with and without incubation with Thrb or bivalirudin for 24 h at 37°C. (**F**) Cell surface proteolysis kinetics tracking the loss of myc tag after the addition of Thrb (5 nM). Data points were fitted to a one-phase exponential decay model. (**G**) Cell surface kinetics tracking the restoration of myc tag after complete linker cleavage with Thrb (200 nM) for 30 min followed by the removal of Thrb. Data points were fitted to a one-phase association model. (**H**) BFP reporter expression kinetics of thrombin-activatable HER2 PAR T cells co-incubated with HER2+ MDA-MB-468 cells after addition of Thrb (200 nM). Data points were fitted to a one-phase association model. (**I**) BFP reporter decay kinetics following removal of Thrb and HER2+ tumor cells. Data points were fitted to a one-phase exponential decay. One-way ANOVA and mean ± SD is depicted, n = 3 biologically independent wells, ****P<0.0001.

To quantify mimotope removal kinetics, we stained HER2 PAR T cells for the upstream Myc tag (**Supplementary Fig. 1D, E**) and observed that linker proteolysis occurred with a half-life of approximately ∼22 minutes (**Fig. 1F** and **Supplementary Fig. 1F**). Considering that cells can internalize their cell surface equivalents at a rate of one to five times per hour^48^, we reasoned that if an extracellular protease was no longer present, cleaved PARs on the cell surface would be replaced by their fully masked counterparts through receptor recycling. To test this, cleaved PAR T cells were transferred to fresh media to remove Thrb, revealing a half-maximal Myc tag reappearance time of ∼8.7 hours (**Fig. 1G and Supplementary Fig. 1G**). We quantified downstream BFP expression kinetics in HER2 PAR T cells continuously co-cultured with HER2-positive MDA-MB-468 cells and Thrb, observing a half-maximal expression time constant of ∼61 hours (**Fig. 1H** and **Supplementary Fig. 1H**) and a half-life of ∼68 hours (**Fig. 1I** and **Supplementary Fig. 1I**).

To assess whether PAR T cells can differentiate on- and off-target proteolysis, we cloned HER2 PARs with LVPRGSG^30,47^, IEFDSG^49,50^, and PLGLAG^30,50,51^ substrate linkers, which are selectively cleaved by Thrb, granzyme B (GzmB) and matrix metalloproteinase (MMP) 9, respectively (**Fig. 2A-C**). All three PAR T cells showed significant BFP upregulation (P<0.0001) when treated with their respective on-target protease but not with off-target proteases. To assess activation selectivity within a cellular mixture, an ∼1:1:1 ratio of PAR T cells displaying PLGLAG, IEFDSG, and LVPRGSG linkers were labeled with either blue (CellTrace^TM^ Blue), green (CellTrace^TM^ CFSE), or red (CellTrace^TM^ Far Red) fluorescent dyes, respectively (**Fig. 2D and Supplementary Fig. 3A-B**). In samples treated with Thrb, we found that approximately 90% of BFP reporter-positive T cells displayed the on-target substrate (LVPRGSG) compared to T cells that displayed either the substrate for GzmB (2.6%) or MMP9 (4.0%) (**Fig. 2E and Supplementary Fig. 3C**). Similarly, treatment of co-cultures with GzmB or MMP9 led to selective reporter expression by cells displaying the respective on-target substrate (80.2% IEFDSG for GzmB and 64.9% PLGLAG for MMP9).

**Figure 2.**
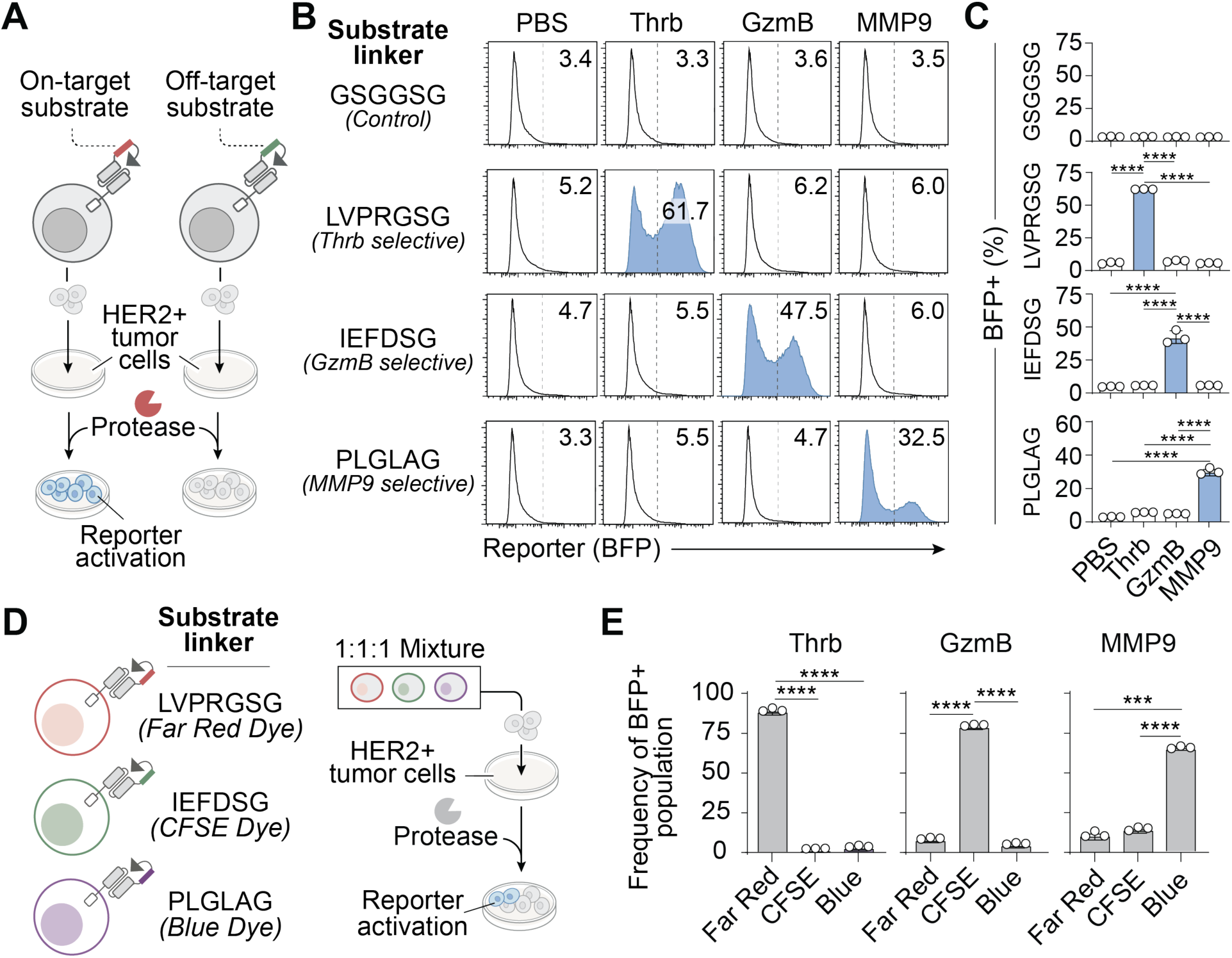
Protease-substrate selectivity drives orthogonal activation of individual HER2 PAR T cells within a three-plex mixture. (**A**) Primary human T cells were engineered to express a HER PAR with substrate linkers LVPRGSG, IEFDSG, or PLGLAG, which are selective for the proteases Thrb, granzyme B (GzmB), or matrix metalloproteinase 9 (MMP9), respectively. The substrate linker GSGGSG was used as a negative control. Each PAR T cell was separately co-incubated with HER2-positive MDA-MB-468 cells with the target or non-target protease (100 nM) at 37°C for 24h. (**B**) Representative flow histogram and (**C**) bar plots of BFP expression frequency show selective activation of PAR T cells after incubation with the target protease, but not with a non-target protease. (**D**) PAR T cells expressing the LVPRGSG, IEFDSG, or PLGLAG substrate linker were labeled with CellTrace^TM^ Far Red, CFSE, or Blue dye, respectively. A three-plex mixture containing approximately equal frequency of each PAR T cell population was co-incubated with either Thrb, GzmB, or MMP9 and HER2-positive MDA-MB-468 cancer cells. (**E**) Bar plots of the frequency distribution of T cells that were positive for CellTrace^TM^ Far Red, CFSE, or Blue dye after gating into BFP-positive PAR T cells from the three-plex mixture, demonstrating orthogonal activation. One-way ANOVA, mean ± SD is depicted, n = 3 biologically independent wells, ***P<0.001, ****P<0.0001.

Together, these results establish that PARs retain protease-substrate selectivity, become antigen-responsive upon unmasking, and revert to their masked state when the activating extracellular protease is no longer present.

### PAR design limits off-tumor T cell activation in antigen-positive healthy tissues

Broad expression of tumor-associated antigens in normal tissues^52^ can lead to on-target/off-tumor activation^53^, as evidenced by dose-limiting toxicities that have largely precluded the clinical success of chimeric antigen receptor (CAR) targets like EGFR^54^ and HER2^55,56^. We postulated that masked PARs could improve spatial specificity and reduce signaling in healthy tissues to enable targeting of broadly expressed antigens. We selected EGFR as it is expressed in most epithelial and vascularized tissues^57^, and cloned primary murine T cells to express either a hEGFR PAR with the proteolytically stable GSGGSG linker or an hEGFR synNotch receptor with the identical scFv. Whereas GSGGSG PAR T cells were insensitive to plate-bound recombinant hEGFR, synNotch T cells exhibited dose-dependent activation (K_D_= 7.1 nM) (**Fig. 3A**). To assess activation *in vivo*, we used a transgenic B-hEGFR mouse in which the EGFR extracellular domain is humanized but otherwise expressed in healthy tissues under its endogenous promoter (liver, lung, kidneys, heart, and stomach; **Fig. 3B, Supplementary Fig. 4A-C**). Following ACT, PAR T cell activation was significantly reduced by >75% in the major organs compared to hEGFR synNotch T cells (**Fig. 3C, D**), confirming PAR masking attenuates antigen-driven activation in healthy tissues.

**Figure 3.**
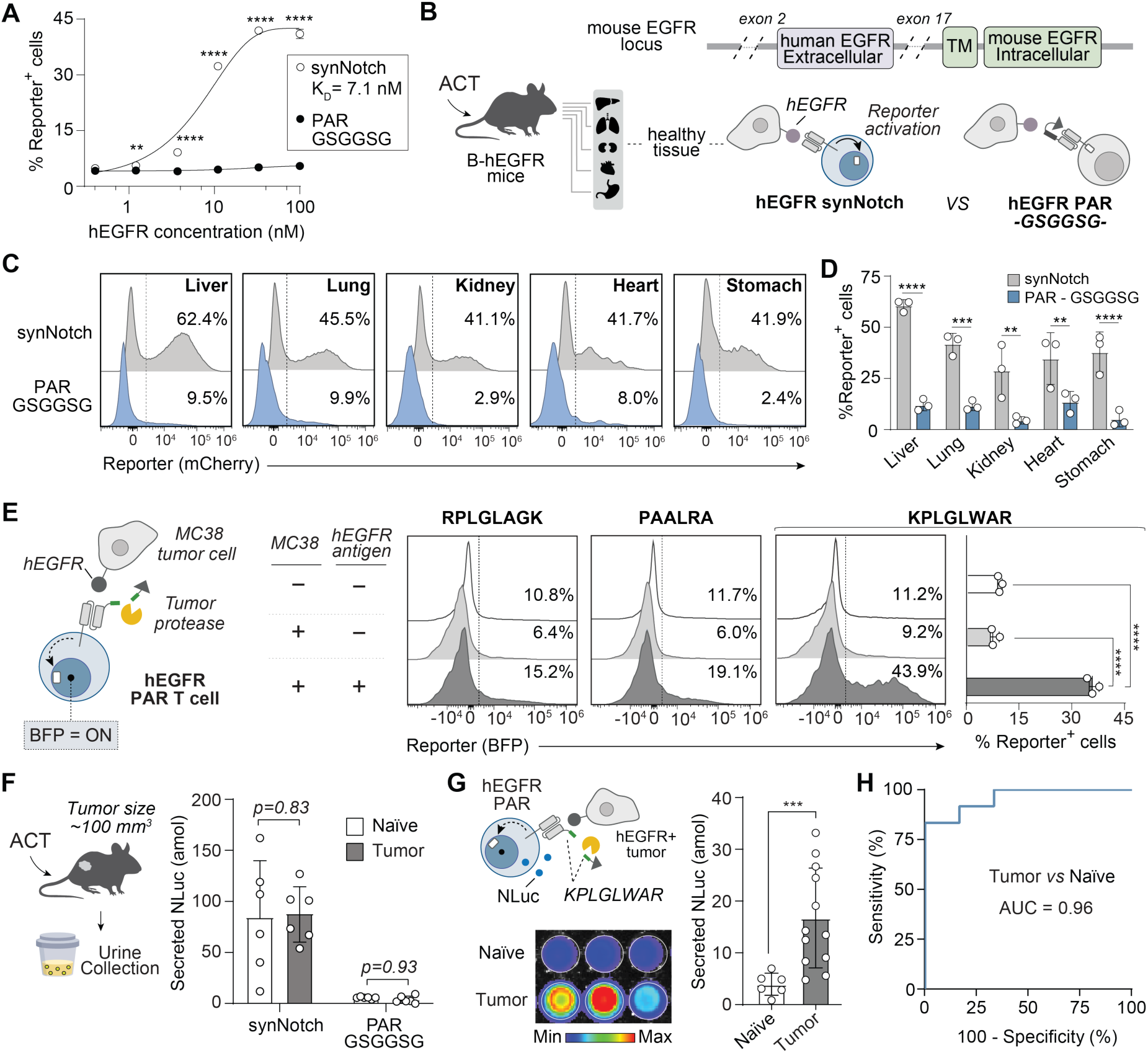
Reducing off-tumor activation allows hEGFR PAR T cells to sensitively detect cancer via a synthetic biomarker secreted in urine. (**A**) Dose-response curve of hEGFR synNotch T cells or hEGFR PAR T cells with control linker (GSGGSG) to plate-bound recombinant human EGFR Fc. (**B**) Schematic of the EGFR locus of B-hEGFR transgenic mice showing replacement of the murine EGFR extracellular domain by its human counterpart. (**C**) Flow cytometry histograms of reporter positive T cells expressing hEGFR synNotch or EGFR PAR presenting GSGGSG control linker isolated from major organs 24 hours after i.v. injection into B-hEGFR mice. Organs were dissociated to single cells, and T cell activation (gated on CD45+CD3+GFP+) was assessed by mCherry expression. (**D**) Bar plots comparing the frequency of mcherry reporter expression across organs. ****P<0.0001, ***P<0.001, **P<0.01, Two-way ANOVA, *n* = 3 biological replicates, error bars depict mean ± SD. (**E**) Representative flow histograms and bar plots showing BFP reporter expression by hEGFR PAR T cells presenting substrate linkers RPLGLAGK, PAALRA, and KPLGLWAR after 24-hour co-incubation at a 1:1 ratio with hEGFR-positive MC38 cancer cells at 37°C. (**F**) hEGFR synNotch or PAR T cells presenting GSGGSG control or (**G**) KPLGLWAR substrate linkers were administered i.v. into hEGFR+ MC38 tumor-bearing mice (∼100 mm^3^) and naïve mice. Urine was collected over a three-hour period one day after injection, and luminescence was quantified using a standard curve of recombinant NLuc. T-test and mean ± SD is depicted, *n* = 6-12 biological replicates, ***P<0.001. (**H**) Receiver operating characteristic (ROC) curves and area under the curve (AUC) values showing diagnostic performance of hEGFR PAR T cell sensors (AUC = 0.96, 95% CI = 0.87-1.00, P = 0.002).

### Murine T cells expressing an MMP-selective PAR detect cancer via secretion of a synthetic urinary biomarker

To evaluate on-tumor activation, we engineered αhEGFR PAR T cells with either RPLGLAGK^30,50,51^, PAALRA^36^, or KPLGLWAR^58^ linkers, which were previously shown to be selective for MMPs. We confirmed that hEGFR PAR T cells displaying each of these linkers were activated upon co-culture with hEGFR-expressing MC38 colorectal cancer cells (**Fig. 3E**), which overexpress MMPs (e.g., MMP1, 2, 9, 10; **Supplementary Fig. 5A**). Among the tested substrates, KPLGLWAR led to the highest PAR activation and was selected for further studies. Using a fluorogenic peptide containing KPLGLWAR flanked with a fluorophore (5(6)-carboxyfluorescein, FAM) and quencher (DABCYL) pair, we confirmed that incubation with several MMPs (MMP1, 7, 12, 13) led to increased fluorescence whereas minimal activation was observed with 12 other proteases (**Supplementary Fig. 5B**), confirming that KPLGLWAR is selectively cleaved by cancer-associated MMPs.

Designing synthetic biomarkers for urinary rather than serum detection can achieve high signal-to-noise ratios, in part because freely filtered reporters are substantially concentrated during tubular water reabsorption^59^. The glomerular filtration size barrier also excludes cells and large serum proteins, creating a simpler matrix for detection and limiting the carryover of components like proteases, opsonins, and phagocytes that reduce reporter stability in circulation. Therefore, we evaluated whether KPLGLWAR PAR T cells engineered to secrete NanoLuc luciferase (NLuc) as a synthetic biomarker, which is rapidly cleared by renal filtration (clearance half-time: 17.7 min; **Supplementary Fig. 6A**), could detect murine tumors. We verified that murine KPLGLWAR PAR T cells released NLuc in co-culture with hEGFR+ MC38 cells, producing 1.7-fold higher luminescence vs. hEGFR– cells (**Supplementary Fig. 6B-C**). We then administered either hEGFR synNotch or GSGGSG PAR T cells to naïve and tumor-bearing B-hEGFR mice (∼100 mm^3^) and confirmed that urinary reporter levels could not distinguish tumor-bearing from naïve animals, with GSGGSG PAR T cells showing significantly reduced overall reporter output (**Fig. 3F** and **Supplementary Fig. 7A-B**). By contrast, PAR T cells displaying KPLGLWAR produced a greater than 4-fold elevation in NLuc reporters in tumor-bearing mice compared to naïve controls (**Fig. 3G**), corresponding to an area under the receiver-operating-characteristic (AUROC) curve of 0.96 (**Fig. 3H**). These results indicate that PARs displaying a tumor-selective substrate can significantly reduce background activation in healthy tissues while using on-tumor activation to drive the release of a synthetic biomarker for detection in urine.

### PAR T cell display enables identification of protease-selective substrates *in vitro*

We next aimed to establish a generalizable pipeline for designing customized PARs for human cancer detection. Protease activity *in vivo* is both tightly regulated^60,61^ and exhibits substrate promiscuity^62^. Existing *in vitro* approaches – such as phage, yeast, and bacteria display – have proved useful for identifying potential protein targets^63^, deciphering substrate cleavage motifs^63–65^ and quantifying enzyme kinetics^66^ but are challenging to implement in the context of living animals to identify protease-cleavable substrates. We therefore sought to evaluate whether PAR T cells displaying a library of substrate linkers could be applied for deep profiling of the substrate repertoire of human cancers *in vivo*.

We expressed a library of HER2 PARs with fully randomized substrate linkers four amino acids in length (4-mers) in primary human T cells, resulting in a unimodal substrate distribution in which >99% of all possible 4-mers (159,978 of 160,000 sequences) were detected by next-generation sequencing (NGS; **Fig. 4A** and **Supplementary Fig. 8**). To confirm the potential for substrate mapping *in vitro*, the library was incubated with 25 proteases across the five major catalytic classes (serine, aspartic, threonine, metallo, and cysteine)^67^ in co-culture with HER2-positive MDA-MB-468 cancer cells. NGS analysis of BFP reporter-positive T cells revealed distinct substrate clusters that were enriched compared to the untreated PAR T cell library for each protease tested (**Fig. 4B** and **Supplementary Fig. 9A**). To address potential sampling bias that could contribute to differences in NGS read counts independent of protease activity, we performed a bootstrapping analysis to simulate sampling of the PAR library across 100 experimental wells (**Supplementary Fig. 9B, C**). This analysis allowed us to down-select to 16,947 substrates with minimal sampling bias while preserving protease-selective substrate clustering (**Fig. 4C** and **Supplementary Fig. 9A**). To further support protease-substrate selectivity, we quantified the frequency of substrates containing the proline-arginine (ProArg) dipeptide within each protease-selective cluster, as Thrb^47^ is known to preferentially cleave after ProArg residues in the P2 and P1 positions. The cluster associated with Thrb showed the highest ProArg enrichment (**Fig. 4D**), consistent with position-specific scoring matrix (PSSM) analysis identifying proline and arginine as the most frequent residues in positions 1–3 and 2–4, respectively, in the thrombin cluster (**Fig. 4E**).

**Figure 4.**
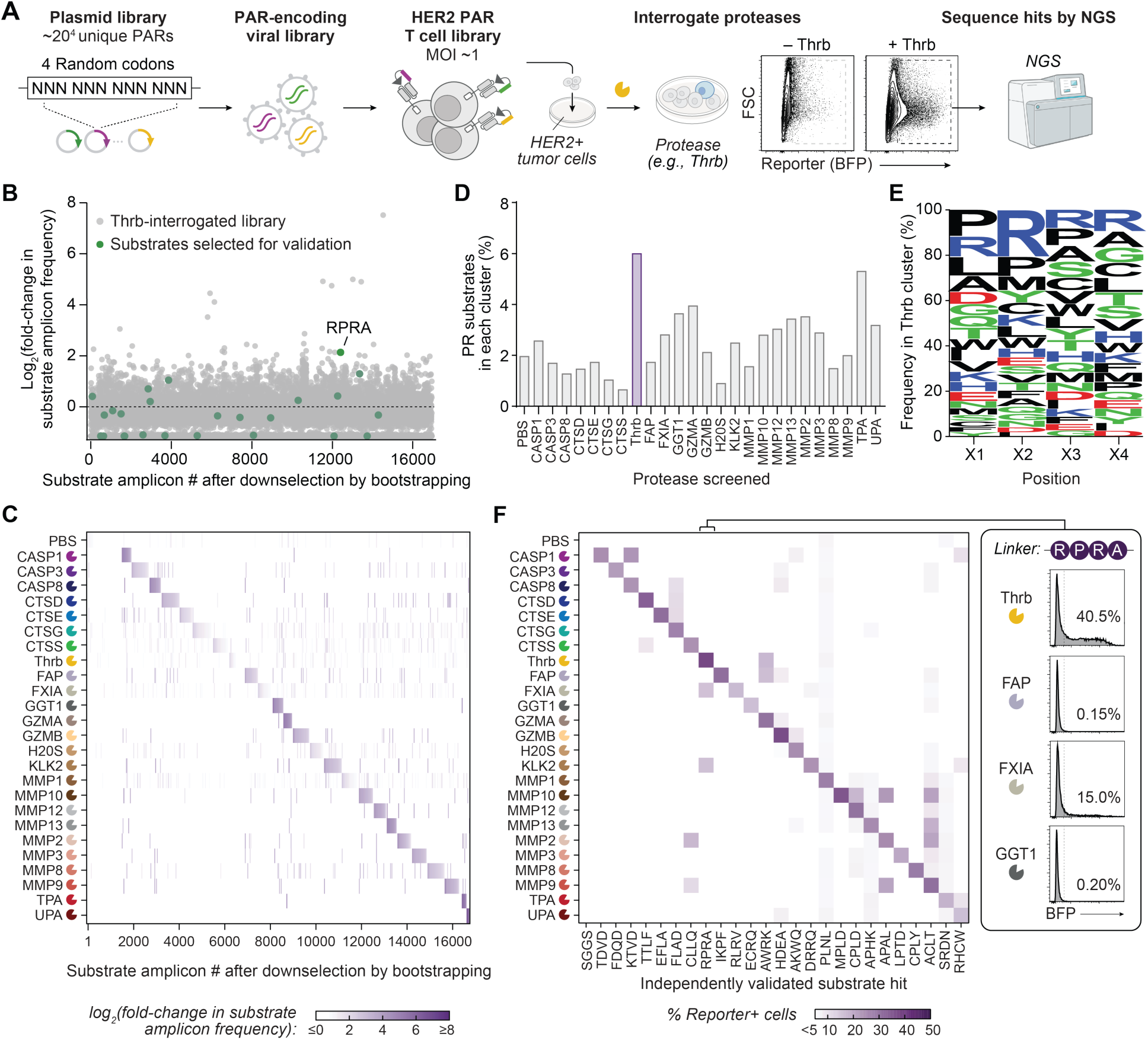
PAR T cell display identifies orthogonal substrates selective for recombinant human proteases *in vitro*. (**A**) Schematic of the *in vitro* PAR T cell display pipeline for screening protease-substrate selectivity. Primary human T cells were engineered by lentiviral transduction to express a library of HER2 PARs that display all possible 4-amino-acid substrate linkers, corresponding to a library diversity of 160,000. The PAR T cell library was co-incubated with one of 25 recombinant proteases and HER2-positive MDA-MB-468 breast cancer cells. Cells were sorted by BFP expression, and their genomic DNA was analyzed by next-generation sequencing (NGS) to identify cleaved substrate sequences. (**B**) Representative dot plot showing the fold-change in amplicon frequency of substrate linkers after treating the PAR T cell library with thrombin. (**C**) Heat map of the log2 fold-change in amplicon frequency for 16,947 substrate linkers tested against 25 proteases, following down-selection from the initial library of approximately 160,000 substrates through bootstrapping. (**D**) Frequency of the dipeptide motif PR appearing in the down-selected set of 16,947 substrates from the respective protease’s cluster. (**E**) Sequence logo generated from position-specific sequence matrix of substrates in thrombin cluster. (**F**) Independent validation of hit sequences from *in vitro* screening using monoclonal PAR T cells. A hit sequence from each cluster was validated by co-incubation of monoclonal HER2 PAR T cells displaying the hit sequence with HER2-positive MDA-MB-468 cells and either the target protease or each of the 24 non-target proteases at 37°C for 24 hours. The heat map shows the average frequency of BFP-positive PAR T cells. n = 3 biologically independent wells.

To experimentally validate that hits discovered from PAR T cell display represent on-target substrates, we engineered 25 distinct monoclonal PAR T cells, each expressing a single enriched substrate selected from one of the 25 protease-substrate clusters (**Supplementary Fig. 10**) and tested their activation by on- and off-target proteases. Notably, all PAR T cells were significantly activated (P<0.0001) by their respective on-target protease (**Fig. 4F** and **Supplementary Fig. 11**), and fourteen showed high selectivity with no detectable activation by any of the other proteases tested (P>0.05). These included substrates cleaved by collagenases (MPLD, APHK, and LPTD for MMP-1, MMP-8, and MMP-13, respectively), cathepsins (TTLF and EFLA for CTSD and CTSE, respectively), and caspases (TDVD and FDQD for CASP1 and CASP3, respectively) despite the redundancy in protease selectivity for native protein substrates within these families^68,69^. These results provide support for the use of PAR T cell display for profiling the substrate repertoire of human proteases *in vitro*.

### PAR T cells selected by *in vivo* display demonstrate tumor-selective activation

We next investigated whether *in vivo* PAR T cell display could be used to screen for tumor-selective substrates in a human xenograft model. Primary human T cells displaying a 4-mer HER2 PAR library were adoptively transferred by intravenous (i.v.) administration into NSG mice bearing either HER2-positive or HER2-negative MDA-MB-468 breast tumors (**Fig. 5A**). Flow cytometric analysis revealed a significant increase in the percentage of BFP-positive T cells isolated from HER2-positive tumors (19.6%) compared with HER2-negative tumors (4.6%) (**Fig. 5B, C** and **Supplementary Fig. 12**). By contrast, PAR T cell activation in the blood, spleen, liver, and lungs – which may arise from ligand-independent activation^70^ – did not differ significantly between tumor-bearing and naïve mice.

**Figure 5.**
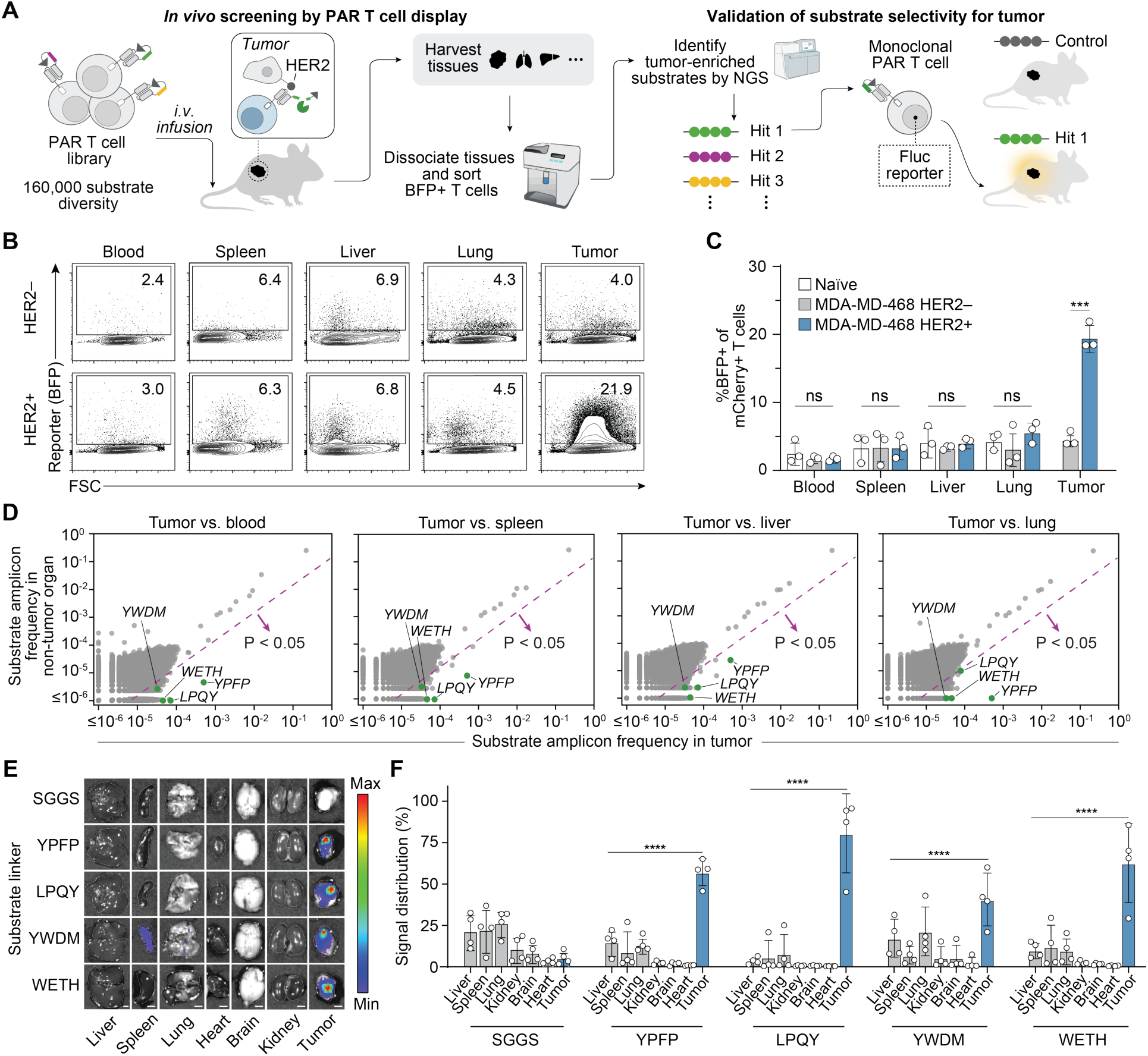
*In vivo* PAR T cell display enables deep profiling of MDA-MB-468 tumors to identify substrates with high tumor selectivity. (**A**) Schematic of *in vivo* PAR T cell display for substrate discovery followed by validation of substrate selectivity in separate cohorts of animals. For discovery, an approximately 62.5-fold representation of a 4-mer library (ten million cells) of HER2 PAR T cells was intravenously (i.v.) injected in NSG mice bearing subcutaneous HER2+ MDA-MB-468 tumors. Twenty-four hours post injection, BFP-positive T cells were isolated from tumor, blood, and non-tumor organs (spleen, liver, lungs) and enriched substrates were identified by NGS. For validation, the top hits were cloned as monoclonal PARs into T cells with a reporter cassette modified to express firefly luciferase (Fluc) to allow validation of tumor selectivity by bioluminescent imaging of tumors and major organs. (**B**) Representative flow plots and (**C**) quantified frequency of BFP expression by PAR T cells in indicated tissues 24 hours after i.v. injection of PAR T cell library to naïve NSG mice or mice bearing HER2-positive or HER2-negative MDA-MB-468 tumors. One-way ANOVA and Tukey post-test and correction, mean ± SD is depicted, n = 3, ***P < 0.001. (**D**) NGS data depicting the frequency of amplicons encoding each substrate linker enriched in the tumor compared to blood, spleen, liver, and lungs. The four highly enriched sequences selected for validation – YPFP, LPQY, YWDM, and WETH – are indicated in green. (**E**) Bioluminescent images of MDA-MB-468 tumors and organs (liver, spleen, lungs, kidneys, brain, heart, and tumor) harvested from mice after the administration of PAR T cells displaying substrate YPFP, LPQY, YWDM, WETH, or SGGS (control). The T cells were modified to include the firefly luciferase (Fluc) gene to report on PAR activation. (**F**) Biodistribution of luminescent signal for PAR T cells displaying each hit sequence or SGGS control. One-way ANOVA and Tukey post-test and correction, ****P < 0.0001, *n* = 4 biological replicates, error bars depict mean ± SD.

NGS analysis of BFP-positive T cells from HER2+ tumors revealed 441 substrate amplicons that were significantly enriched compared to the blood, spleen, liver, and lungs (**Fig. 5D**). From these, we selected 4 highly enriched sequences (YPFP, LPQY, WETH, and YWDM) for validation, using monoclonal PAR T cells cloned with a UAS-Firefly luciferase (Fluc) reporter gene to enable quantitative assessment of local tissue activation by bioluminescent imaging (**Supplementary Fig. 13A**). Adoptive transfer of each PAR T cell into separate mouse cohorts resulted in significantly higher luminescent signals in tumors compared with the liver, spleen, lungs, kidneys, brain, and heart (P<0.0001 for YPFP, LPQY, and YWDM; P<0.001 for WETH) (**Fig. 5E, F** and **Supplementary Fig. 13B**). By contrast, PAR T cells displaying a SGGS control linker did not lead to measurable increases in tumor signal above background levels.

Because LPQY showed the highest tumor-to-organ signal ratio, we advanced this sequence for further validation to confirm that PAR T cell activation is mediated by tumor-selective protease activity. Given that MDA-MB-468 tumors express MMPs at high levels *in vivo* (MMP1, 8, 9, 10, 12, and 13) (**Supplementary Fig. 14A**), we reasoned that tumor-associated MMP activity contributed to LPQY cleavage. Using a FAM/DABCYL fluorogenic peptide, we found that LPQY was selectively cleaved by 8 MMPs, while exhibiting minimal activity against 17 additional proteases (**Supplementary Fig. 14B-C**). *In vivo*, LPQY HER2 PAR T cell activation was significantly reduced in HER2-positive tumors pre-treated with an intratumoral injection of marimastat, a broad spectrum MMP inhibitor, compared to untreated tumors. Additionally, luminescent signals were also reduced in HER2-negative tumors regardless of marimastat treatment, confirming that LPQY PAR T cell activation is both protease- and antigen-dependent (**Supplementary Fig. 15**). These results support the use of *in vivo* PAR T cell display for deep profiling of the tumor substrate repertoire to identify tumor-bespoke PARs.

### OncoSCOUT achieves a lower tumor detection limit than protein and ctDNA benchmarks

We next assembled all components to create a customized OncoSCOUT test using HER2 PAR T cells with the LPQY substrate and engineered to release nLuc as a synthetic urinary biomarker for human cancer detection (**Fig. 6A**). In mice bearing ∼100mm^3^ MDA-MB-468 tumors, OncoSCOUT produced significantly higher synthetic urinary biomarkers one day after infusion compared to PAR T cell controls displaying a SGGS substrate linker (**Fig. 6B**). Daily 3-hour urine collections over three consecutive days revealed that urinary signal peaked on day 1 and decreased over the subsequent days (**Fig. 6C**). Consequently, day 1 was selected for all subsequent comparisons.

**Figure 6.**
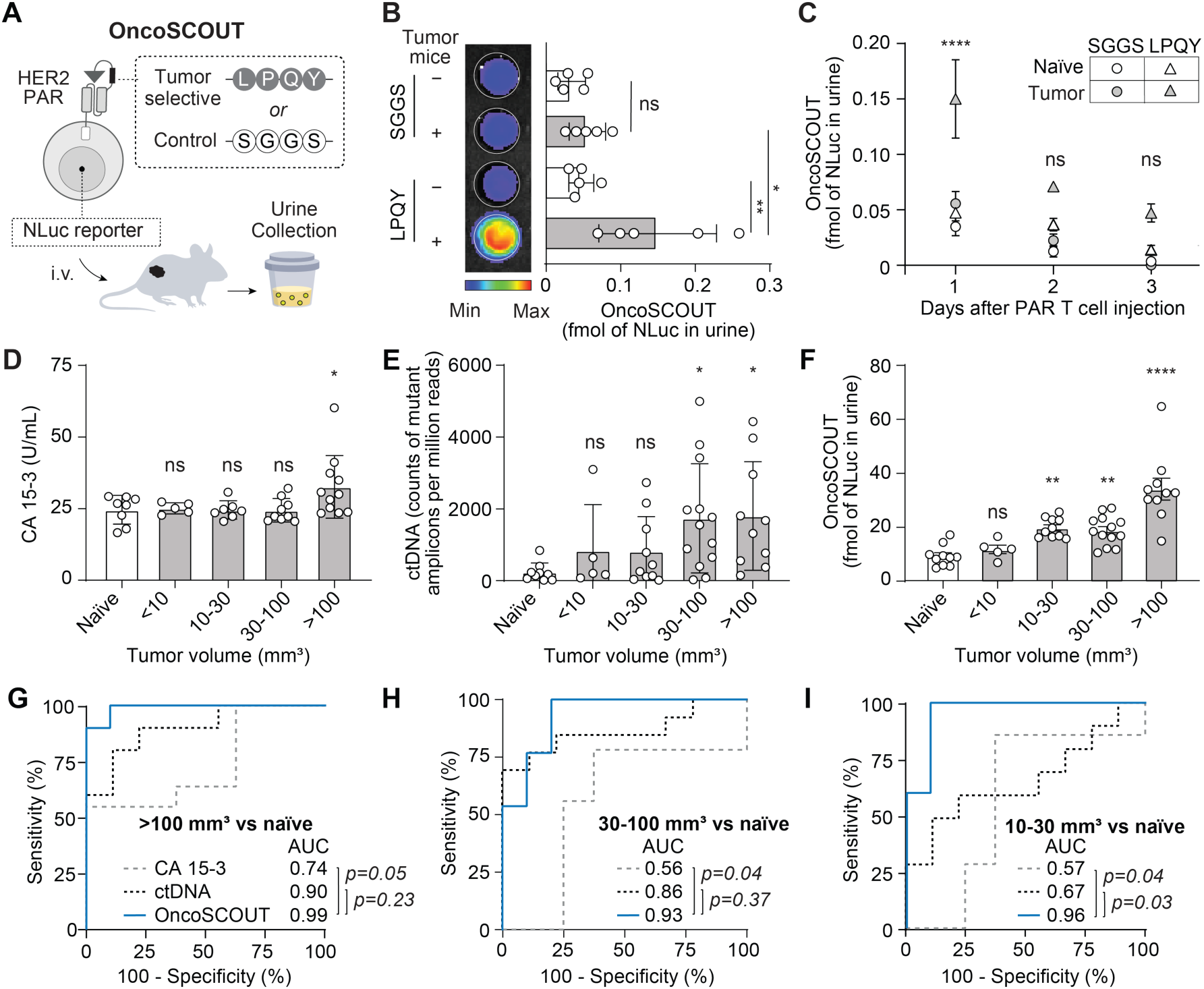
OncoSCOUT detects smaller tumors with greater sensitivity and specificity compared to CA 15-3 and ctDNA. (**A**) Schematic of customized OncoSCOUT assay comprising HER2 PAR T cells with substrate LPQY encoding Nluc as a synthetic urinary biomarker. (**B**) Luminescent quantification of NLuc levels in urine samples collected from HER2+ MDA-MB-468 tumor-bearing and naïve mice. Urine was collected over a 3-hour window one day after injection or (**C**) on days 2 and 3. Urinary luminescence was quantified using a standard curve generated from recombinant NLuc on the IVIS Spectrum CT system. Two-way ANOVA, P values compare urine samples from naïve and tumor-bearing mice. (**D**) Serum CA 15-3 levels across increasing tumor size intervals compared to naïve controls quantified by ELISA. (**E**) Frequency of mutant DNA amplicons in plasma from naïve and tumor-bearing mice. Plasma was collected from the same cohort used for OncoSCOUT evaluation. Targeted sequencing was performed using a custom targeted gene panel. (**F**) Synthetic urinary biomarker levels from OncoSCOUT as a function of increasing tumor size. Urine was collected over a 3-hour window one day after T cell administration, and luminescence was quantified using the IVIS Spectrum CT system using a recombinant NLuc standard curve. ****P<0.0001, **P<0.01, *P<0.05, one-way ANOVA, *n* = 5-13 biological replicates, mean ± SD is depicted. (**G**) ROC curves and corresponding AUC values evaluating the classification performance of CA 15-3, ctDNA and OncoSCOUT tests for tumor sizes greater than 100 mm^3^, (**H**) between 30 and 100 mm^3^, and (**I**) 10 to 30 mm^3^. DeLong test was used for the statistical comparison of the AUCs.

To benchmark the tumor-size detection limit, we compared OncoSCOUT with CA 15-3 – an FDA-approved protein biomarker used to monitor breast tumor progression and recurrence^71,72^ – which is expressed by MDA-MB-468 tumor cells (**Supplementary Fig. 16**) and a multiplex ctDNA assay. We collected serum and EDTA-plasma samples from separate cohorts of mice for CA 15-3 and ctDNA, respectively, to minimize cfDNA background from clotting-induced lysis of leukocytes. The ctDNA cohort received an OncoSCOUT infusion, and after one day, plasma and urine samples were collected at the same time to directly compare detection accuracy. Across tumors of increasing volume (<10, 10–30, 30-100 and >100 mm^3^, **Supplementary Fig. 17**), serum CA 15-3 levels were significantly elevated only in mice with tumors exceeding 100 mm^3^ (**Fig. 6D** and **Supplementary Fig. 18A**), whereas levels in mice with smaller tumors were statistically identical to naïve mice and remained within the physiological range of 0–30 U/mL^73^. For comparison with ctDNA, we used a targeted amplicon sequencing panel covering over 2,900 known hotspots in human cancers to identify a fingerprint of 20 mutations across 17 oncogenes and tumor suppressor genes in the gDNA of MDA-MB-468 breast cancer cells that were not present in healthy human T cells (**Supplementary Table 1**). We observed a significant increase in the total frequency of these 20 mutant amplicons in mice with tumors ranging from 30–100 mm^3^ in volume, but not in those with tumors smaller than 30 mm^3^ (**Fig. 6E** and **Supplementary Fig. 18B**).

By contrast, OncoSCOUT produced synthetic urinary biomarker signals that detected tumors identifiable by CA 15-3 (>100 mm^3^) and ctDNA (>30 mm^3^), while also distinguishing tumors as small as 10–30 mm^3^ that the other tests did not significantly detect (**Fig. 6F** and **Supplementary Fig. 18C**). To evaluate diagnostic sensitivity and specificity, receiver-operating-characteristic (ROC) curve analysis across tumor size bins (**Supplementary Fig. 18D–F**) demonstrated that OncoSCOUT significantly outperformed CA 15-3 for tumors >100 mm^3^ based on the area under the curve (AUC) (0.99 vs. 0.74) while showing comparable tumor discrimination to ctDNA (0.99 vs. 0.90) (**Fig. 6G**). A similar trend was observed for tumors 30–100 mm^3^ (**Fig. 6H**), with OncoSCOUT significantly outperforming CA 15-3 (0.93 vs. 0.56) and maintaining comparable tumor discrimination to ctDNA (0.93 vs. 0.86). In the 10–30 mm^3^ size range (**Fig. 6I**), OncoSCOUT showed the greatest improvement, achieving a significantly higher AUC of 0.96 compared with 0.57 for CA 15-3 and 0.67 for ctDNA. These results indicate that OncoSCOUT can achieve lower cancer detection thresholds than CA 15-3 or a 20-plex ctDNA assay, with significantly improved sensitivity and specificity.

## Discussion

The motivation to develop synthetic biomarkers for earlier detection is driven by mathematical^74,75^ and experimental evidence^76,77^ indicating that endogenous biomarkers have inherent physiological barriers that limit their sensitivity for detecting early-stage disease. Integrating concepts from masked prodrugs^46,78,79^ and synNotch T cells^41,42^, we describe the design of masked PARs that enhance tumor-selective activation by requiring both extracellular protease activity and antigen-dependent signaling. We report a generalizable *in vivo* substrate-screening approach via PAR T cell display to customize OncoSCOUT for the detection of human breast cancer xenografts via a synthetic urinary biomarker, achieving discrimination of established tumors as small as 10–30mm^3^ with significantly improved sensitivity and specificity compared to existing cancer biomarkers. Our study used peptide mimotopes previously reported for cetuximab and trastuzumab to design EGFR and HER2 PARs. This strategy can be extended to additional scFvs lacking characterized peptide mimotopes through high-throughput discovery methods such as yeast display of degenerate peptide libraries^80^ or tiled linear peptide epitopes^81^. We designed PAR T cells to display substrates 4-amino acids in length to enable >10-fold library representation *in vivo*, which requires the adoptive transfer of at least 1.6 million T cells. A limitation of scaling to larger libraries (e.g., 6- or 8-mer) with comparable fold representation is the need to pool samples from multiple mice or to conduct multi-round screens in which enriched 4-mers are progressively extended and diversified at each stage. By comparison, phage, yeast and bacterial libraries can reach library diversities of ∼10^9^. However, yeast and bacterial libraries have primarily been used for *in vitro* substrate mapping due to rapid immune reactivity and clearance in living animals. Phage libraries can be administered for *in vivo* selection with minimal immediate immune response, but they depend on passive tissue targeting and lack the molecular specificity afforded by the PAR architecture. In addition, most prior work has focused on identifying tissue-binding peptides rather than protease-cleavable sequences, which require tailored motifs such as cryptic cleavage sites^82^ or cell-penetrating peptides^83^.

Most advanced cancers carry thousands of somatic mutations^84^, and their detection through ctDNA is driving progress in personalized drug selection^85^, response monitoring^86^, and minimal residual disease assessment^87^. In the setting of early detection, however, tumors smaller than 1 cm^3^ are predicted by mathematical modeling to produce <2 genomic equivalents of DNA in a 10–15 mL blood draw^74,75^, resulting in mutant allele frequencies below 0.01%^88,89^. At these levels, ctDNA can be obscured or surpassed by variant alleles present in normal cfDNA arising from processes like clonal hematopoiesis^90,91^. In our study, OncoSCOUT benchmarked favorably against targeted sequencing of known ctDNA mutations in a xenograft model with tumors ranging from ∼10–100mm^3^, where somatic alterations in normal cfDNA were readily identifiable by their murine origin and did not confound the detection of human-derived ctDNA. However, a limitation of our study is that the relationship between tumor burden and biomarker levels in mice compared to humans is not fully understood, and mathematical modeling will likely be required to predict how sensitivity scales allometrically – such as with differences in body size, blood volume, tumor size and urine production – across species^59,92^. Additionally, our comparisons were limited to protein biomarker analysis and targeted ctDNA sequencing. Future work should evaluate OncoSCOUT against other approaches, such as methylation profiling, which will require large datasets and machine learning-based classifiers to accurately interpret and classify signals^93,94^.

Several additional limitations of this study merit further investigation. The high cost of *ex vivo* cell manufacturing and adoptive transfer currently precludes routine cancer screening with engineered cells; however, non-integrating *in vivo* cell engineering approaches – such as the use of lipid nanoparticles for transient gene delivery^95,96^ – could potentially allow OncoSCOUT to be developed as an off-the-shelf product. Furthermore, our observation that synthetic biomarker signals could distinguish tumor-bearing animals within one day of adoptive transfer suggests that short-lived allogeneic cells^35,97^ could potentially be used for tumor detection, which offers the opportunity for scalable manufacturing and lower cost of goods. Notably, although there are recognized differences in the protease proteome across species, approximately 500 murine proteases are orthologues of human enzymes^98^, and protease-activatable probes, including the FDA-approved pegulicianine, have demonstrated comparable efficacy in both human xenograft models and patients^23,99^. Nonetheless, further studies are needed to characterize protease expression and the corresponding degradome in primary human tumor samples to design PARs that best enable cancer early detection. Finally, an initial clinical entry point could prioritize recurrence surveillance or monitoring of treatment response, particularly in cancer patients receiving adoptive cell therapy. In all settings, clinical imaging is expected to play an important role in identifying the tissue-of-origin to confirm synthetic biomarker levels and could potentially be supported by engineering PAR T cells with genetically inducible MRI, PET, or ultrasound reporters^100,101^.

In summary, our results support the use of PAR T cells for deep profiling of the cancer substrate repertoire to customize early detection of cancer via synthetic urinary biomarkers.

## Materials and Methods

### Animal models

Female NSG mice (6- to 12-week-old), male and female C57BL/6J, male and female B-hEGFR mice (5- to 12-week-old) were used at the outset of all experiments. NSG mice were bred in-house using breeding pairs purchased from The Jackson Laboratory. C57BL/6J mice were purchased from The Jackson Laboratory. B-hEGFR mice expressing human EGFR were bred in-house using breeding pairs purchased from Biocytogen. In B-hEGFR mice, exons 2–17 of the mouse EGFR gene encoding the extracellular domain were replaced with the human counterparts, while the transmembrane, cytoplasmic, promoter, 5′ signal peptide, and 3′UTR regions were retained. Chimeric EGFR expression is driven by the endogenous mouse EGFR promoter, with native mouse EGFR transcription and translation disrupted. All animal procedures were approved by the Georgia Tech Institutional Animal Care and Use Committee (protocol no. A100190, A100191, A100193 and A100677). All authors complied with relevant ethical regulations while conducting this study.

### Cell lines

Human embryonic kidney 293T and MDA-MB-468 cell lines were obtained from the American Type Culture Collection. MC38 cell line was a gift from the National Cancer Institute and D. Vignali, University of Pittsburgh). The cells were maintained in Dulbecco’s Modified Eagle’s Medium (DMEM, Corning) supplemented with 10% fetal bovine serum (FBS, Sigma-Aldrich) and 1% penicillin/streptomycin (Gibco). HER2-positive MDA-MB-468 cells were transduced by retrovirus with full-length human HER2 and sorted by fluorescence-activated cell sorting (FACS) to >95% purity. EGFR-positive MC38 cells were transduced by lentivirus with truncated human EGFR and FACS sorted to >97% purity.

### Primary human T cell isolation from healthy donors

Peripheral blood mononuclear cells (PBMCs) were obtained from healthy donors (IRB no. H20288) by venipuncture and isolated by density gradient using Lymphocyte Separation Media (Corning). CD3+ T cells were enriched from PBMCs using EasySep Human CD3 Cell Isolation kits (Stem Cell Technologies) according to the manufacturer’s instructions. Isolated CD3+ T cells were cultured in human T cell media consisting of X-VIVO 10 (Lonza), 5% human AB serum (Valley Biomedical), 10 mM *N*-acetyl L-cysteine (Sigma-Aldrich), and 55 µM β-mercaptoethanol (Sigma-Aldrich) supplemented with 50 units per ml human IL-2 (Corning). All cells were cultured at 37°C in 5% CO_2_.

### Primary murine T cell isolation

Mouse T cells (mTCs) were isolated from healthy C57BL/6J mouse spleens using the same protocol of our previous work. CD3+ T cells were enriched from splenocytes using EasySep mouse T Cell Isolation kits (Stem Cell Technologies) according to the manufacturer’s instructions. Isolated T cells were then stimulated with mouse CD3/CD28 dynabead (Thermo fisher) with a 1:1 ratio and cultured in mouse T cell media (mTCM) consisting of RPMI-1640 (HyClone) with 10% Fetal Bovine Serum (FBS, Sigma-Aldrich), 1× MEM nonessential amino acids and 1 mM sodium pyruvate (Corning), 55 µM β-mercaptoethanol (Sigma-Aldrich), supplementing with 100 units per ml human IL-2 (Corning). Cells were cultured at 37°C in 5% CO_2_ and maintained at 1×10^6^ cells/mL.

### Plasmid construction

Anti-HER2 synNotch receptors and response elements were obtained from Addgene# 85424 and 79130, respectively. The DNA sequence encoding a human CD8a signal peptide (MALPVTALLLPLALLLHAARP), myc-tag (EQKLISEEDL), peptide mimotope (LLGPYELWELSH), and protease substrate flanked by GS linkers was codon-optimized and cloned into the N-terminus of the scFv in the anti-HER2 synNotch-Gal4VP64 construct. Anti-EGFR was cetuximab-derived scFv and cloned into pMKO.1 retroviral vector which included synNotch receptors and response elements. The DNA sequence encoding mouse CD8a signal peptide (MASPLTRFLSLNLLLLGESIILGSGEA), myc-tag, EGFR peptide mimotope (QGQSGQCISPRGCPDGPYVMY), non-cleavable sequence (GSGGSG), thrombin cleavable substrate (LVPRGSG) or MC38 tumor specific substrate (RPLGLAGK, PAALRA, KPLGLWAR) were codon-optimized for murine and cloned into the EGFR synNotch construct. All cloning was performed using amplification by polymerase chain reaction with Phusion DNA Polymerase (New England Biolabs) and restriction enzyme digestion. Plasmids were verified by Sanger sequencing (Eurofins Genomics), whole plasmid sequencing (Primordium Labs), and/or next-generation sequencing (Admera Health) prior to use. Plasmid DNA was purified using the E.Z.N.A.® Endo Free Plasmid Maxi Kit (Omega Bio-Tek). For generations of plasmid libraries for substrate screening, an oligo library of 160,000 sequences encoding the fully degenerate 4-mer substrate library was synthesized (Twist Bioscience) and cloned into the masked anti-HER2 synNotch receptor plasmid.

### Lentiviral production and human T cell transduction

Lentivirus was produced by co-transfection of lentiviral expression plasmids with psPAX2 and pMD2.G using TransIT-LT1 transfection reagent (Mirus Bio) and HEK293T cells. Viral supernatant was collected after 48 hours, concentrated using PEG-it Virus Precipitation Solution (System Biosciences) following the manufacturer’s protocol, and stored at −80°C until use. Activated human T cells were co-transduced by mixing two viral constructs encoding the PAR (Multiplicity of infection (MOI)=1) and the synNotch-activated reporter (MOI=5). Concentrated lentivirus was used to transduce T cells by addition to a 24-well suspension culture plate coated with retronectin (Takara) according to the manufacturer’s instructions, followed by centrifugation at 1,200×g for 90 minutes at 37°C. Activated human T cells were added to each well in human T cell media supplemented with 100 units/mL of human IL-2, and the plate was spun at 1,200×g for 60 minutes at 37°C. Cells were incubated on the virus-coated plate for 24 hours before expansion, and transduction efficiency was evaluated by flow cytometry analysis of cells stained with anti-myc tag antibodies and cells expressing mCherry. Seven days after activation, T cells were supplemented with Dynabeads at a 1:1 bead to cell ratio. The beads were removed on day 9. Cells were maintained at a concentration of 7×10^5^ to 2×10^6^ cells/mL until day 10-14 for use in downstream assays. To minimize manufacturing bias of library production, a minimum 10-fold representation of the library was maintained at each step—plasmid amplification, lentivirus production, T cell transduction, and maintenance.

### Retroviral production and mouse T cell transduction

Retrovirus was produced by co-transfection with pCL-Eco and pMKO.1 retroviral vectors encoding EGFR PAR or Luminescence reporter in HEK293T cells using TransIT-293. Cell supernatant was collected 24 and 48 hours later and combined for filtering through a 0.22 µm rapid-flow filter unit (Nalgene). The retrovirus was store at –80 °C until use. Activated mouse T cells were co-transduced by mixing two retrovirus encoding the hEGFR PAR and the synNotch-activated luminescent reporters. The mixture of 1 mL retrovirus and 4 ng/mL polybrene with 1e6 T cells were added to a 24-well suspension plate pre-coated with retronectin (Takara), followed by centrifugation at 2000 ×g at 32 °C for 2 h. And 1 mL of fresh mTCM supplemented with IL-2 (100 units/ml) were added on the top of each well after 4 h of centrifugation. The transduced T cells were cultured in the 24-well plate overnight and transferred to the centrifugation tubes and spined down at 1000 ×g at RT for 5 min to remove the virus, following by resuspension in mTCM with IL-2 and seeded to a flask at a concentration of 1e6 cells/mL for expansion. T cell transduction was checked after 24 and 48 hours by flow cytometry for expression of myc-tag and mCherry (or eGFP). Dynabeads were removed 48 hours post transduction. The transduced cells were then either sorted for co-expression of myc+ and mCherry+ (or eGFP+) and maintained at 1e6 cells/mL for further *in vitro* studies or injected to mice for *in vivo* studies.

### In vitro PAR T cell activation

Target cancer cells were seeded in a 96-well plate overnight. PAR T cells were co-incubated with seeded cancer cells with or without target protease at a 1:1 ratio at 37°C. The cultures were analyzed for reporter expression after 24 h with a BD Fortessa (BD Biosciences; FACSDiva v8 software) or Aurora (Cytek Biosciences; SpectroFlo v3.2.1 software) flow cytometer. All flow cytometry analysis was performed with FlowJo software.

### hEGFR *synNotch and* hEGFR PAR T cell activation in B-hEGFR mice model

PAR T cells were generated by co-transducing mouse CD3⁺ T cells with retroviral constructs encoding EGFR synNotch or PAR, along with mCherry reporters. Transduced T cells were intravenously injected into B-hEGFR mice. Twenty-four hours after injection, the mice were euthanized, and the liver, lungs, kidneys, heart, and stomach were harvested. T cells were isolated from each organ and stained with Live/Dead NIR (Thermo Fisher), αCD45-PE-Cy7, and αCD3-BV750, each at a 1:50 dilution from stock concentrations. Stained cells were analyzed on an Aurora flow cytometer (Cytek Biosciences) using SpectroFlo v3.2.1 software.

### Synthesis of fluorogenic peptide substrate

Protease substrate peptides with 5(6)-carboxyfluorescein (FAM) fluorophore and N-[4-(4-dimethylamino)phenylazo]benzoic acid (DABCYL) quencher (FAM-[Substrate]-(K-DABCYL)-NH2) were synthesized in-house using the Liberty Blue peptide synthesizer (CEM). The peptide synthesis scale used was 0.025 mmol, and low-loading rink amide resin (CEM) was used. Amino acids (Chem-Impex) were resuspended in dimethylformamide (0.2 M), as were all synthesis buffers. Activator buffer used was diisopropylcarbodiimide (DIC; Sigma) (0.25 M) and the activator base buffer was Oxyma (0.25 M; CEM), while the deprotection buffer was piperidine (20% v/v; Sigma). After synthesis, peptides were cleaved off resin using with a mixture of 92.5% trifluoroacetic acid, 2.5% water, 2.5% triisopropylsilane, and 2.5% 3,6-dioxa-1,8-octane-dithiol for 30-45 minutes at 41°C using Razor (CEM). After cleavage, peptides were precipitated in ice-cold diethyl ether and vacuum-dried overnight. Peptides were validated by liquid chromatography-mass spectrometry (1260 Infinity II HPLC and single quadrupole LC/MSD, Agilent). Purity of all peptides was >80%.

### Fluorogenic substrate cleavage assay

Fluorogenic peptides (5 µM) were mixed with recombinant proteases (50 nM, diluted according to manufacturer’s buffer) or tissue lysate and incubated at 37°C. Fluorescence was measured by Cytation 5 plate reader (Biotek; Gen5 software). Protease concentrations were 50 nM unless otherwise specified. For preparing tissue lysate, tumor and organs were isolated and dissociated using FastPrep-24 homogenizer (MP Biomedicals) with beads (Lysing Matrix D, MP Biomedicals) in tissue protein extraction buffer (T-PER, ThermoFisher). Homogenate was isolated by centrifugation at 14,000xg for 5 minutes.

### Circulation half-life and urine clearance of recombinant NLuc

Recombinant NanoLuc (NLuc) (200 fmol) was administered intravenously to naïve NSG mice. At designated time points post-administration, blood and urine were collected from separate cohorts of mice. Serum was isolated by centrifugation. Samples were mixed with the Nano-Glo® Live Cell Assay System (Promega) and immediately analyzed. Luminescence was quantified using a standard curve generated with recombinant NLuc.

### *PAR T cell sensor for tumor detection* via urinary synthetic biomarkers

PAR T cells were injected i.v. into tumor-bearing mice and naïve mice. Urine was collected for a three-hour window one day after injection or longitudinally for up to 7 days Luminescence was quantified using a standard curve of recombinant NLuc.

### Mapping human protease-substrate repertoire

HER2 PAR T cell library was generated by co-transducing 10 million CD3+ T cells with a HER2 PAR lentiviral library encoding 160,000 substrates sequences (MOI 1) and a lentivirus encoding the synNotch-activated BFP reporter (MOI 5). 2 million FACS-sorted cells were coincubated with one million HER2 expressing MDA-MB-468 cells with or without target protease (100 nM) at 37°C for 24 hours. After incubation for 24 h, activated PAR cells were sorted (FACSAria, BD Biosciences) on BFP expression and pelleted at 1000×g for 3 min before genomic DNA (gDNA) isolation.

### Mapping substrate repertoire in breast tumor model

Ten million FACS-sorted T cells expressing the PAR library were intravenously injected into naïve mice or mice bearing HER2-negative or HER2-positive MDA-MB-468 tumors (∼100 mm^3^ in size). Twenty-four hours after injection, mice were euthanized, and the tumor, spleen, blood, liver, and lungs were collected to isolate T cells. The tumors were minced and digested for 30 min in RPMI with 0.1 mg/mL DNase (Roche) and 0.2 mg/mL collagenase P (Roche) at 37°C. Digested tumors were passed over a 75 µm cell strainer before cells were collected by centrifugation, treated with red blood cell (RBC) lysis buffer (Biolegend), washed with PBS, and resuspended in FACS buffer (1x DPBS, 2% FBS, 1 mM EDTA, 25 mM HEPES). Spleens were gently dissociated using frosted glass slides before splenocytes were centrifuged at 1000xg for 5 min, resuspended in RBC lysis buffer for 5 min at 4°C, washed with PBS, and resuspended in FACS buffer. For blood, 500 µL was collected prior to euthanasia in an EDTA-coated tube. Blood samples were mixed with 5 mL of red blood cell lysis for 5 min at 4°C, washed with PBS, and resuspended in FACS buffer. Livers were forced through a 75 µm cell strainer using the barrel of 5 mL syringe and centrifuged at 500xg for 10 min. The pellets were suspended in 8 mL of 40% Percoll (in RPMI) and overlaid on 5 mL of a 67% Percoll fraction. Percoll gradient separation was performed by centrifugation at 840×g for 20 min at room temperature. The lymphocytes were collected at the interface, washed, and resuspended in FACS buffer. The lungs were minced and digested for 30 min in RPMI with 0.1 mg/mL DNase (Roche) and 0.2 mg/mL collagenase D (Roche) at 37°C. The digested lungs were passed over a 75 µm cell strainer, followed by centrifugation at 500xg for 5 min. The cells were then treated with red blood cell lysis buffer (Biolegend), washed with PBS, and resuspended in FACS buffer. The isolated cells from each organ were stained with αCD3 (300406) at 1:100 dilution from stock concentrations. Stained cells were analyzed and sorted for BFP expression using a FACSAria (BD Biosciences; FACSDiva software).

### Genomic DNA purification and NGS library preparation

The gDNA of sorted BFP+ T cells was extracted using QIAprep Spin Miniprep Kit according to manufacturer’s protocol. At least two million unsorted library cells (>10 times the diversity of the library) were collected from culture and prepared as an input sample for each screen. The library was amplified with the following primers, which contain overhangs encoding the binding sites and adaptors: F5’ TGCTTGGTCCTTATGAGCTG and R5’ GTGAGGCTCTGCAGGTTATTG. Library preparation and NGS sequencing (50 million paired end reads) were performed by Admera Health using an Illumina NovaSeq.

### Extraction of amplicon counts from NGS paired end reads

Raw FASTQ sequencing files containing paired end reads were analyzed using custom code on R. Briefly, the code used the ShortRead package to iteratively input forward and reverse reads, filter out misaligned and low-quality (Phred<30) paired-end reads, and translate reads into amino acid sequences. Sequences were further filtered by excluding sequences with mismatches in the 9 residues upstream or downstream of the random 4-mer region and sequences in which the 4-mer region encoded a stop codon. The final list of sequences was tabulated into amplicon counts and exported to Microsoft Excel for hit selection.

### Analysis and hit selection for in vitro screening

To generate the initial heat map without bootstrapping, we converted amplicon counts for samples generated from each protease into frequencies, which were normalized to fold-changes using the corresponding frequencies in the original library. Substrates were clustered based on the protease that generated the highest fold-change value for each sequence. Heat maps were generated from the log2 fold-change values using the Seaborn package on Python. Bootstrapping analysis was performed using custom code on R. Hits were selected by rank-ordering the substrates in each protease’s cluster based on highest fold-change in amplicon frequency and choosing one of the top 20 substrates.

### Analysis and hit selection for in vivo screening

We converted amplicon counts for samples generated from each organ (tumor, blood, spleen, liver, and lungs) into frequencies. Substrates that were not detected in the tumor were excluded from further analysis. For all remaining substrates, we calculated the ratio of the frequency in the tumor to the frequency in each of the non-tumor organs and determined which substrates had tumor-to-organ ratios greater than 1.96 standard deviations above the mean, corresponding to a P-value of 0.05. Tumor-selective substrates were defined as substrates with significant tumor selectivity (P < 0.05) compared to all four non-tumor organs.

### In vivo bioluminescence imaging to validated PAR encoded hit substrates

PAR T cells were generated by co-transducing CD3+ T cells with lentiviral constructs for HER2 PAR encoding hit substrate sequences (YPFP, LPQY, YWDM and WETH) and Firefly luciferase (Fluc) reporters. Five million PAR T cells were intravenously injected into mice bearing HER2-expressing MDA-MB-468 tumors (100 mm3). Twenty-four hours after injection, mice were euthanized, and the tumor, spleen, blood, liver, lungs, kidneys, brain, and heart were harvested. The organs were immersed in D-luciferase (Gold Biotechnology, 90 µg/mL) solution for 30 min. Fluc activity was measured using an IVIS Spectrum CT (PerkinElmer).

### Plasma preparation and ctDNA mutation analysis

Blood was collected via terminal cheek bleed into EDTA-coated tubes and maintained on ice until processing. Plasma was separated by two-step centrifugation: 1,600 × *g* for 10 min at 4 °C, followed by 14,000 × *g* for 10 min at 4 °C to remove residual debris. The clarified plasma was stored at −80 °C until further use. ctDNA was extracted using the Mag-Bind® cfDNA Kit (Omega Bio-tek) according to the manufacturer’s protocol. Library preparation and next-generation sequencing (NGS) were performed by Admera Health using the OncoZoom® Cancer Panel (Paragon Genomics). For identifying the mutational fingerprint, sequencing data from gDNA of MDA-MB-468 cells was processed by filtering out low-quality reads (Phred score < 30), and forward read amplicons that were not found in healthy human T cells and were distinct from the Genome Reference Consortium Human Build 38 sequence were identified as mutant amplicons. For analysis of plasma samples, the total read count was quantified as all reads with Phred score ≥ 30. Reads for each mutant amplicon were further processed by removing misaligned paired-end reads. Subsequently, the ratio of reads across the 20 identified mutant amplicons to the total read count was quantified for each mouse.

### Statistical analysis

Appropriate statistical analyses were performed using GraphPad Prism (*P < 0.05, ** P < 0.01, *** P < 0.001, **** P < 0.0001). Central values represent mean and error bars depict SD. At least 3 replicates were used for all statistical analyses. Flow cytometry data were analyzed using FlowJo X (FlowJo, LLC). Power analyses were performed using G*Power 3.1 (HHUD). Ex vivo and in vivo luminescence data were collected and analyzed with Living Image 4.4.5 (PerkinElmer). Figures were designed in Adobe Illustrator.

## Data Availability

The main data supporting the results in this study are available within the paper and its Supplementary Information. Other data generated and analyzed during the study are available from the corresponding author on reasonable request.

## Acknowledgements

This work was supported by the National Cancer Institute (NCI) of the National Institutes of Health (NIH) under grant numbers DP1CA280832 and DP2CA280622, and by the Advanced Research Projects Agency for Health (ARPA-H) under award number AY2AX0000006. A.S. was supported by the National Science Foundation (NSF) Graduate Research Fellowships Program under grant number DGE-2039655. A.Z. was supported by the Ruth L. Kirschstein National Research Service Award (NRSA) for Individual Predoctoral Fellows (F31) under grant number CA271803. This work was performed in part at the Georgia Tech Institute for Electronics and Nanotechnology, a member of the National Nanotechnology Coordinated Infrastructure, which is supported by the National Science Foundation under grant number ECCS-1542174. This content is solely the responsibility of the authors and does not necessarily represent the official views of the National Institutes of Health or the Advanced Research Projects Agency for Health. We thank the staff at Georgia Tech’s Cellular Analysis and Cytometry Core and Department of Animal Resources for their assistance in performing our studies.

## Author Contributions

H.P. and G.A.K. conceived the idea. H.P., Y.C., A.S., A.Z., L.G., Q.D.M., H.J.L., L.R., J.B., P.Q., and G.A.K. designed experiments and interpreted results. H.P., Y.C., A.S., A.Z., L.G., Q.D.M., H.J.L., L.R., J.Y., S.A.S., S.Z., and M.S.G. synthesized materials and carried out the experiments. H.P., A.S., Y.C., and G.A.K. wrote the manuscript.

