## Supplementary Information for "Programming T cells for Early Cancer Detection with Customized Protease-Activatable Receptors"

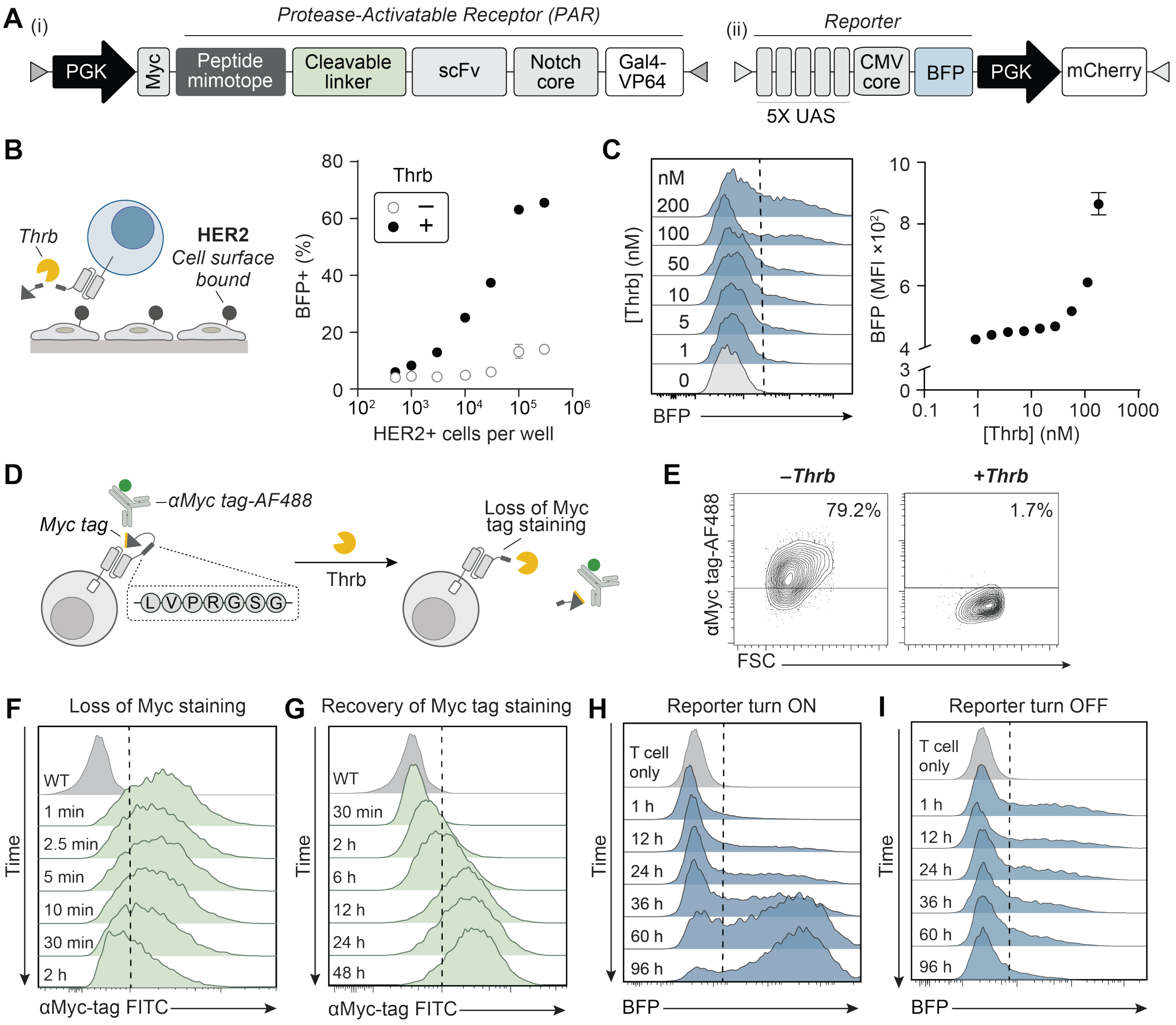


**Supplementary Figure 1. Activation of HER2 PAR T cells is antigen- and protease-dose dependent and fully reversible upon protease removal.** (**A**) Schematic of gene circuits for (i) masked Protease-Activatable Receptor (PAR) and (ii) 5x UAS-CMV promoter upstream of a blue fluorescent protein (BFP) reporter. (**B**) Dose response of HER2 PAR T cells engineered with Thrb-activatable linker (LVPRGSG) upon addition of Thrb (200 nM) with varying levels of HER2 expressed by MDA-MB-468 cancer cells and (**C**) the concentration of Thrb in co-culture. (**D**) Schematic of myc tag staining to monitor substrate cleavage. (**E**) Flow plots of primary human T cells engineered with Thrb-activatable PAR show a decrease in myc tag staining upon proteolysis for 30 min at 37°C.

(**F**) Histogram show proteolysis kinetics measured by loss of staining with αmyc tag antibody upon Thrb addition (5 nM); (**G**) after complete cleavage is reached (200 nM Thrb for 30 min), removal of Thrb induces recovery of αmyc tag staining indicating substrate turnover. (**H**) Histogram show BFP reporter expression kinetics upon Thrb addition (200 nM); after 24 hours incubation, (**I**) removal of Thrb and HER2+ tumor cells. Mean ± SD is depicted, n = 3 biologically independent wells.

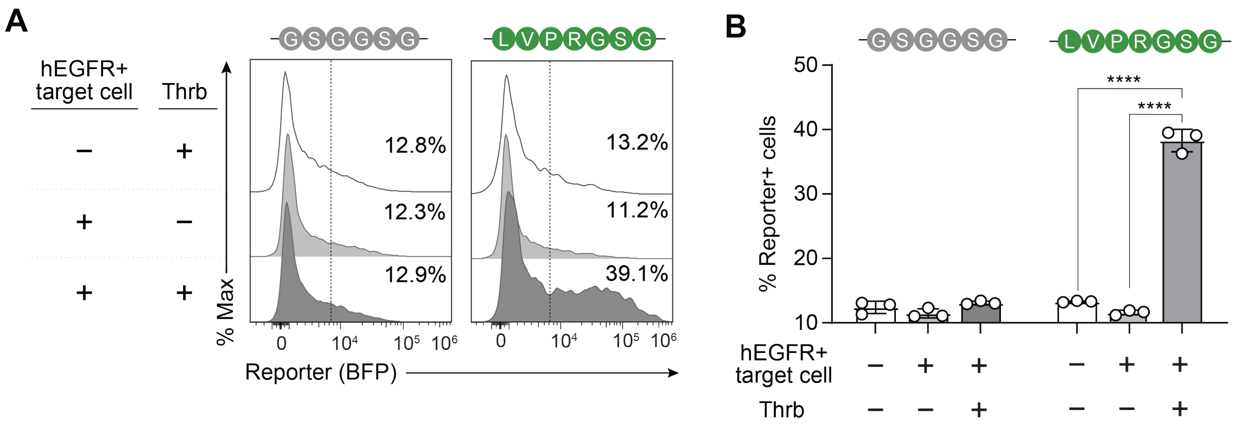


**Supplementary Figure 2. Activation of hEGFR PAR T cells requires an extracellular protease and hEGFR antigen. (A)** Flow histogram and (**B**) bar plots of primary murine T cells engineered with thrombin-activatable (LVPRGSG) or control (GSGGSG) hEGFR PARs show that expression of the BFP reporter requires both thrombin (Thrb) and hEGFR-positive MC38 colon cancer cells (+ + condition).

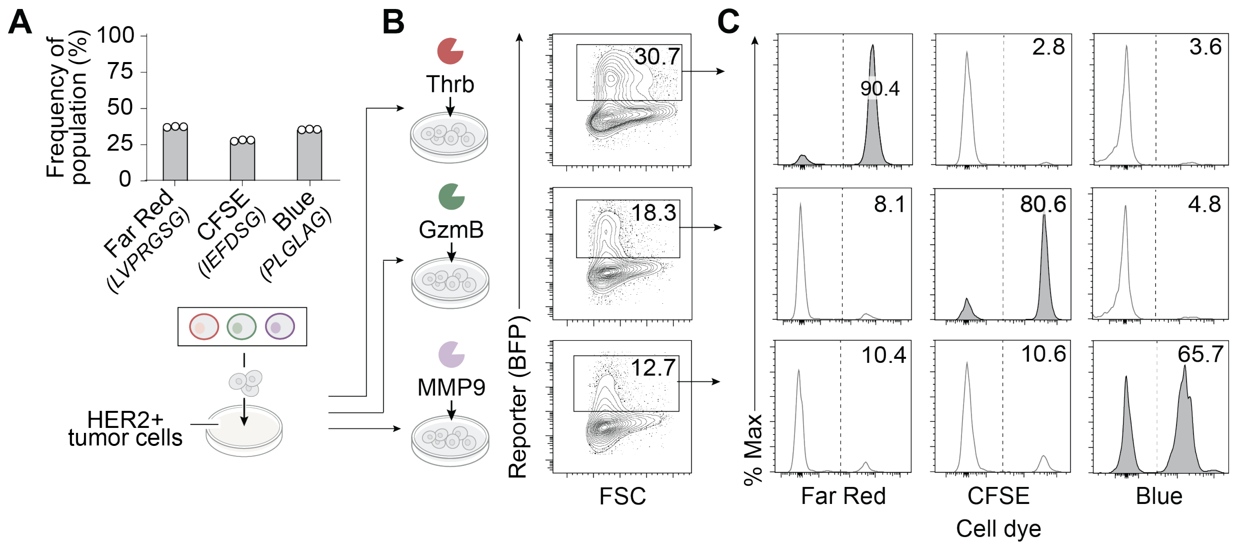


**Supplementary Figure 3. PAR T cell activation in a mixed population depends on the substrate specificity of extracellular proteases.** (**A**) HER2 PAR T cells expressing LVPRGSG, IEFDSG, or PLGLAG substrate linkers were labeled with CellTrace$^{TM}$ Far Red, CFSE, or Blue dyes, respectively. These cells were combined at ~1:1:1 ratio. The bar plot depicts the cell distribution quantified by flow cytometry. (**B**) A polyclonal mixture of the three cell populations was co-incubated with the indicated protease and HER2+ MDA-MB-468 cancer cells, resulting in BFP expression in a fraction of the cells as indicated in the flow plots. (**C**) Gating on BFP expressing cells, the histograms show the distribution of dye-labeled PAR T cell clones in the BFP-positive population (Far Red 630/661, CFSE 492/517, Blue 355/410). n = 3 biologically independent wells.

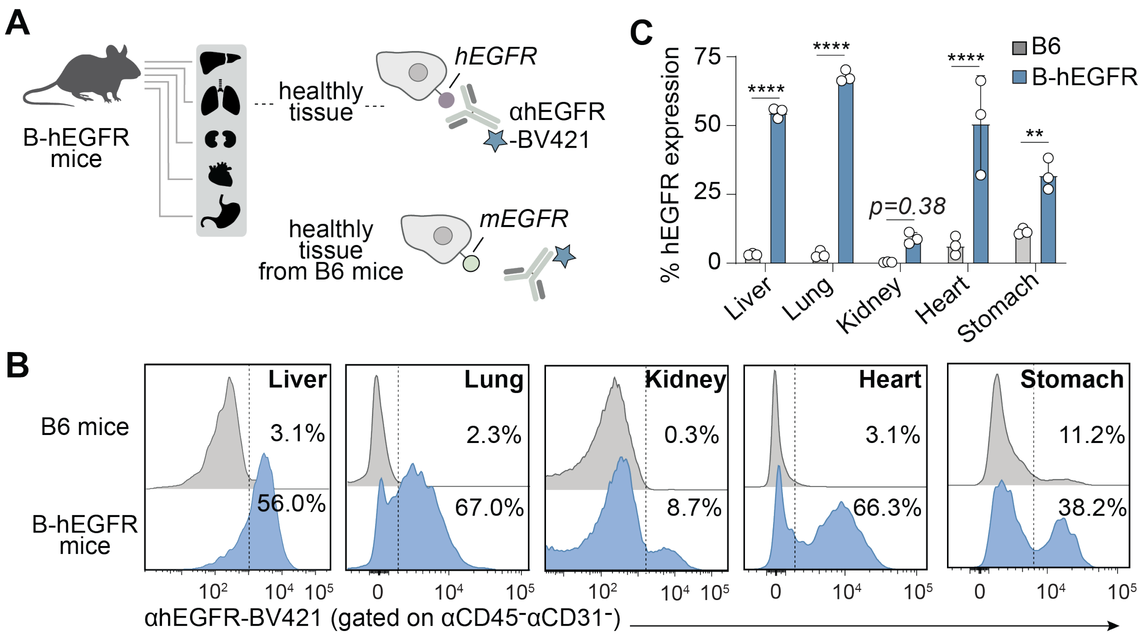


**Supplementary Figure 4. Off-tumor hEGFR expression in healthy tissues of B-hEGFR transgenic mice induces reporter activation in hEGFR synNotch T cells.** (**A**) Schematic of the B-hEGFR transgenic mouse model, in which the mouse EGFR extracellular domain was replaced with the human counterpart. hEGFR-expressing tissues engage αhEGFR scFv, triggering Gal4-VP64 release and reporter expression from the UAS promoter in synNotch receptors. (**B**) Representative flow plots and (**C**) bar graphs of human EGFR expression in tissues including liver, lung, kidneys, heart and stomach in B-hEGFR and C57BL/6 (B6) mice. ****P<0.0001, ***P<0.001, Two-way ANOVA, *n* = 3 biological replicates, error bars depict mean ± SD.

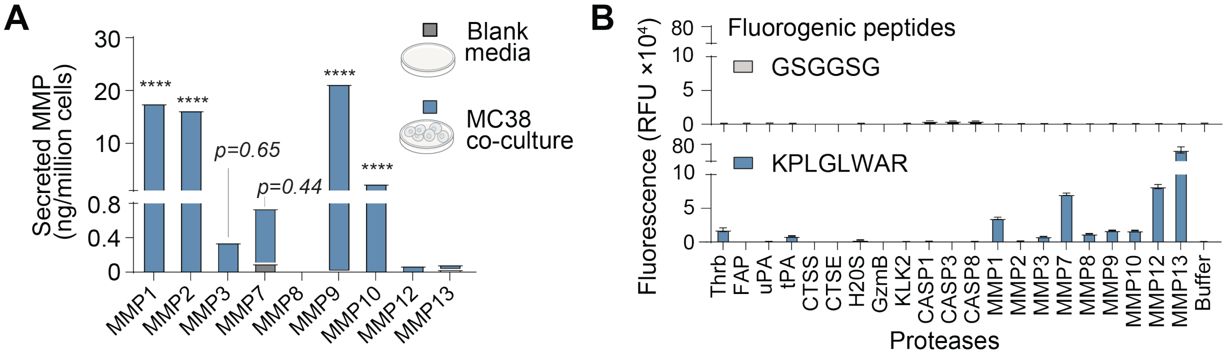


**Supplementary Figure 5. The substrate KPLGLWAR is primarily cleaved by matrix metalloproteinases.** (**A**) MMP protease expression in MC38 cancer cells. One million cancer cells were cultured at 37ºC for 48 hours. The supernatant was collected and analyzed by ELISA. ****P<0.0001, Two-way ANOVA, *n* = 3 biological replicates, mean ± SD is depicted. (**B**) The fluorescence of samples containing either the KPLGLWAR or GSGGSG (control) fluorogenic peptides after incubation with recombinant proteases (50 nM) for 30 min at 37°C.

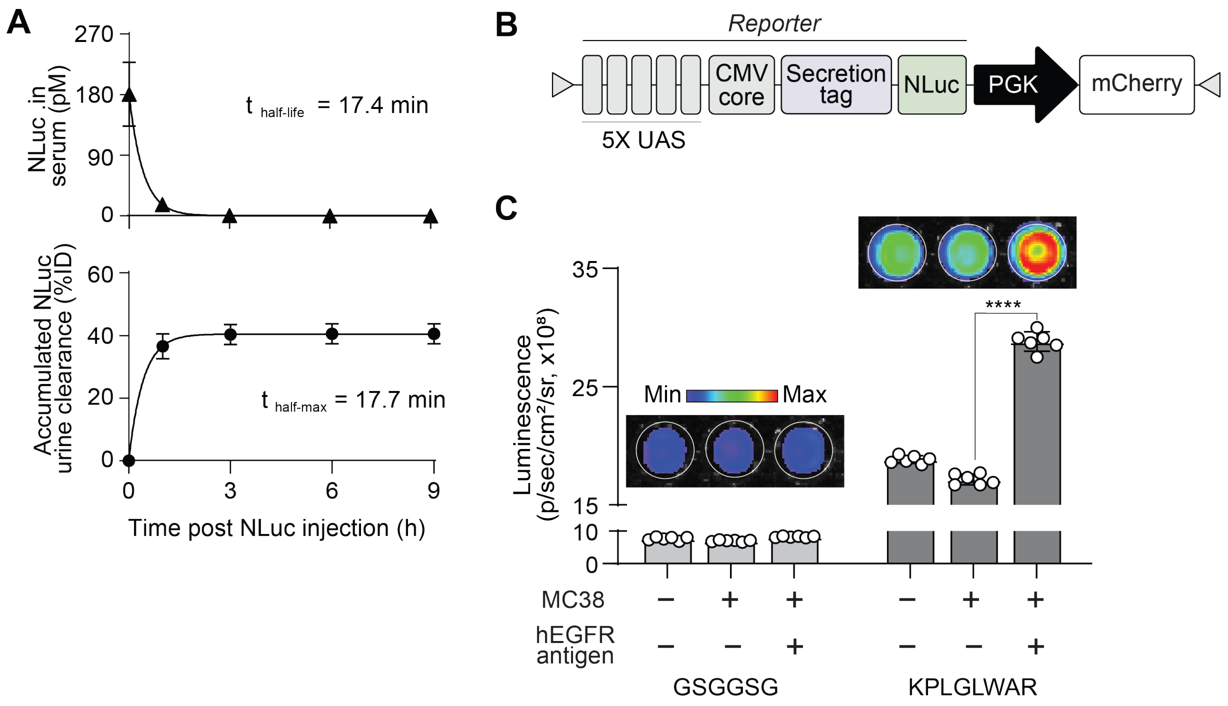


**Supplementary Figure 6. Recombinant NLuc is rapidly cleared into urine *in vivo* and can be produced by hEGFR PAR T cells in vitro.** (**A**) Measurement of NLuc half-life in serum and urine after i.v. administration of 200 fmol recombinant protein in mice. Data was fitted to a one-phase exponential decay or association model. $n = 3$ biological replicates, error bars depict mean ± SD. (**B**) Gene map of reporter plasmid encoding NLuc modified with a secretion tag. (**C**) Quantification of luminescence in the supernatant of 24-hour co-cultures at 37°C containing primary mouse T cells expressing hEGFR PAR with KPLGLWAR or GSGGS control linkers and $hEGFR^{+}$ or $hEGFR^{-}$ MC38 cancer cells. One-way ANOVA, **** P<0.0001, $n = 6$ biological replicates, error bars depict mean ± SD.

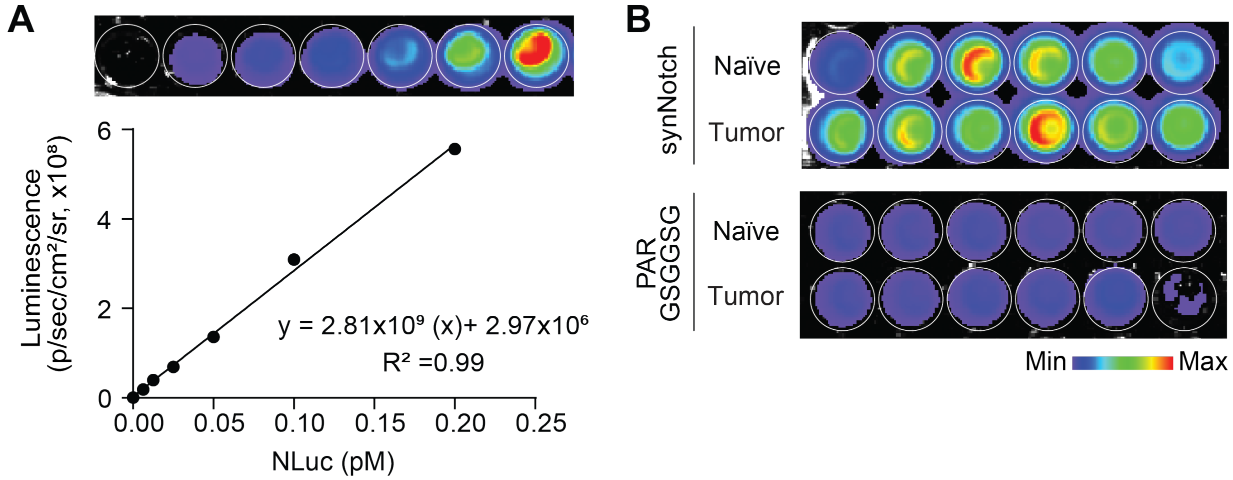


**Supplementary Figure 7. Quantification of luminescence in urine from mice bearing hEGFR⁺ MC38 tumors after administration of hEGFR synNotch or control hEGFR PAR T cells.** (**A**) Standard curve of recombinant NLuc quantified with the IVIS Spectrum CT system. (**B**) IVIS images of urine samples collected from mice bearing hEGFR⁺ MC38 tumors over a three-hour period one day after injection of synNotch or control PAR T cells.

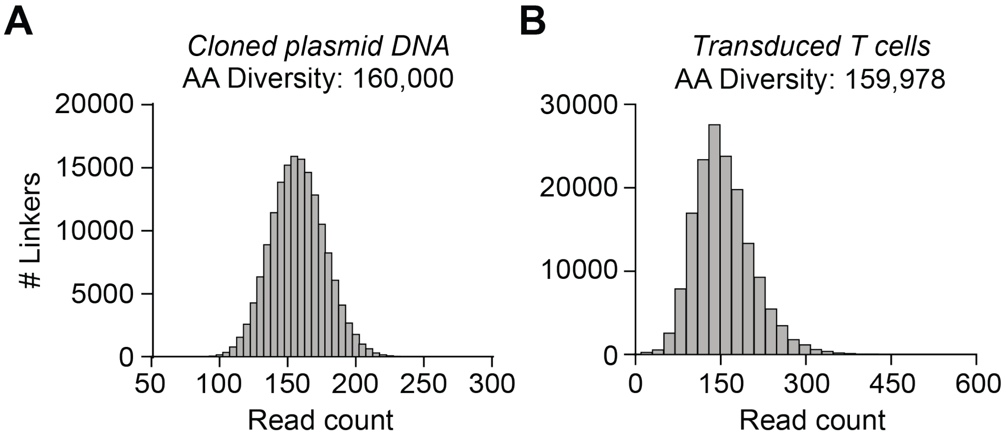


**Supplementary Figure 8. Substrate distribution of PAR T cell library.** T cells were engineered by lentiviral transduction to express a library of HER2 PARs that display 4-amino-acid substrate linkers, corresponding to a theoretical library diversity of 160,000. Histograms depict frequency distribution of substrates in (A) PAR plasmid DNA library and (B) T cells encoded PAR library.

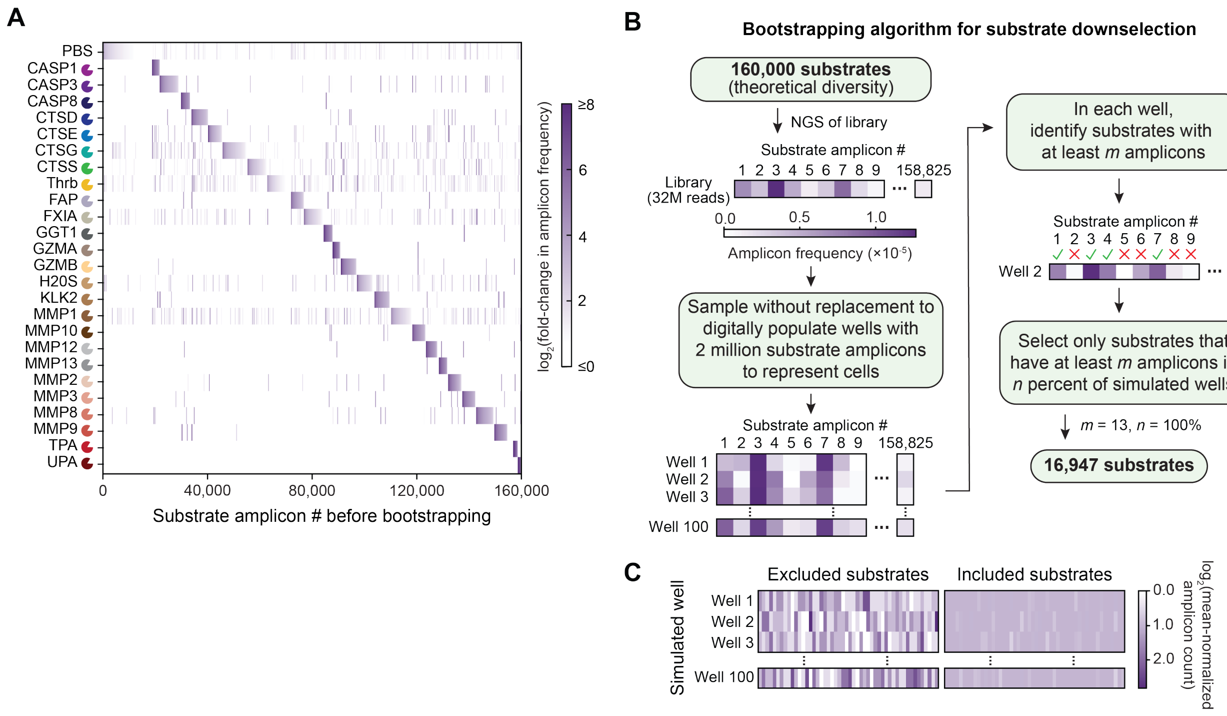


**Supplementary Figure 9. Bootstrapping algorithm identifies substrates with minimal sampling bias.** (**A**) Heat map summarizing log2 of fold-change in amplicon frequency of all ~160,000 substrates after PAR T cell display against 25 proteases. (**B**) Schematic of algorithm to bootstrap wells containing 2 million substrate linkers from the non-enriched library and filter out sequences predicted to have high sampling bias. (**C**) Heat map demonstrates that bootstrapping algorithm includes substrates predicted to have lower variance across wells from random sampling and excludes substrates predicted to have higher variance.

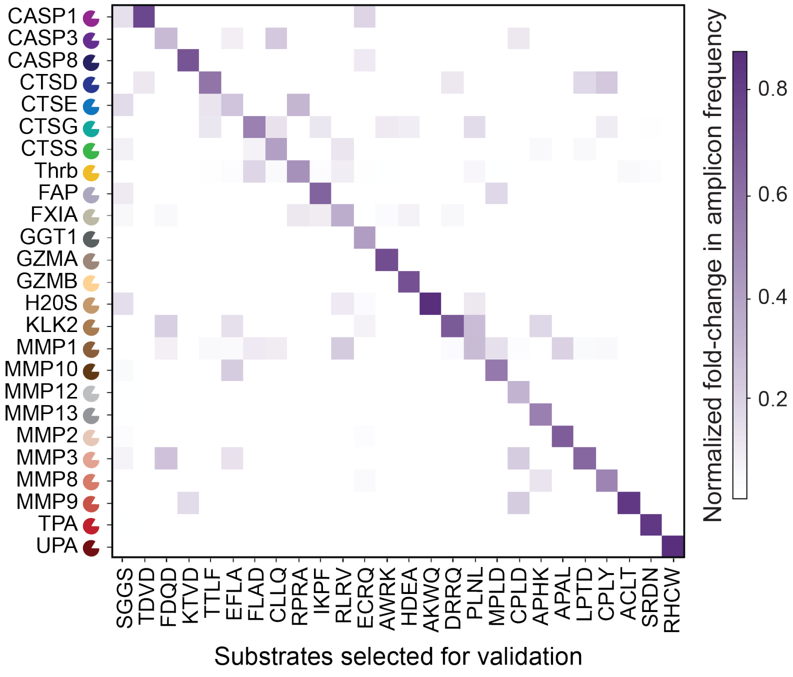


**Supplementary Figure 10. Heat map of substrate sequences identified by NGS analysis and selected for downstream validation.** Heat map of fold-change in amplicon frequency of SGGS control linker and 25 selected hit substrates against 25 proteases. Fold-change values were normalized for each substrate linker.

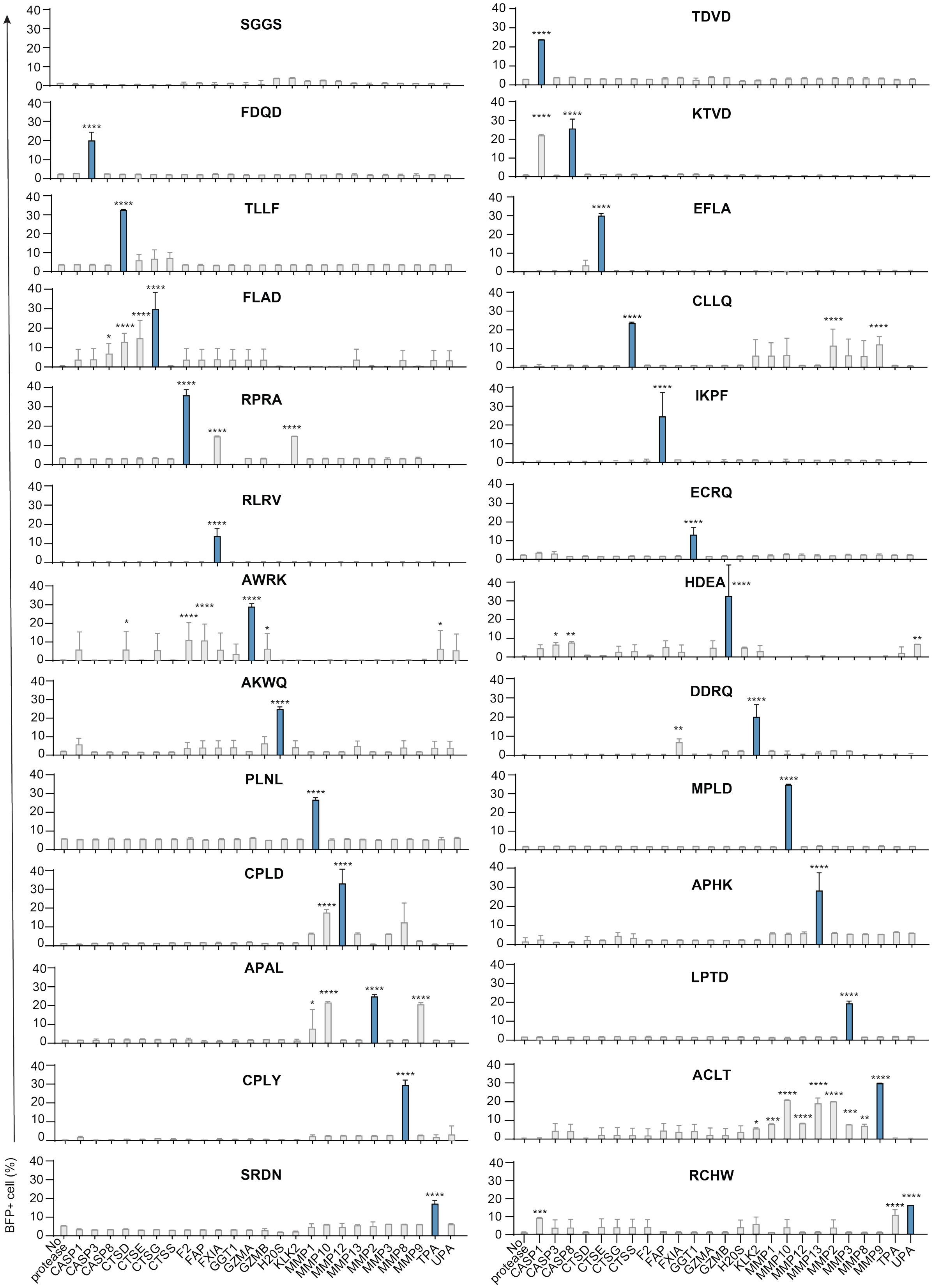
SGGS
TDVD
FDQD
KTVD
TLLF
EFLA
FLAD
CLLQ
RPRA
IKPF
RLRV
ECRQ
AWRK
HDEA
AKWQ
DDRQ
PLNL
MPLD
CPLD
APHK
APAL
LPTD
CPLY
ACLT
SRDN
RCHW
BFP+ cell (%)
0
10
20
30
40
No protease
CASP1
CASP3
CASP8
CTSD
CTSE
CTSG
CTSS
F2
FAP
FXIA
GGT1
GZMA
GZMB
H20S
KLK2
MMP1
MMP10
MMP12
MMP13
MMP2
MMP3
MMP8
MMP9
TPA
UPA

**Supplementary Figure 11. Quantification of reporter expression in monoclonal PAR T cells validating hit sequences from *in vitro* screening.** Hit sequences from each protease cluster were validated by co-culturing monoclonal HER2 PAR T cells displaying individual hit sequences with MDA-MB-468 HER2+ cells and the corresponding target protease for 24 hours at 37ºC. The bar plot shows the average PAR T cell BFP reporter expression. *P<0.05, **P<0.01, ***P<0.001, ****P<0.0001, two-way ANOVA comparing with no protease samples. n = 3 biologically independent wells.

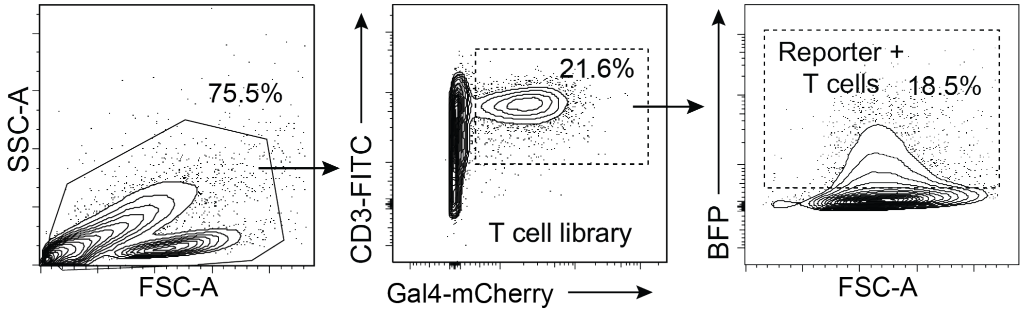


**Supplementary Figure 12. Gating strategy to sort reporter-activated PAR T cells from tumor for *in vivo* PAR T cell display screening.**

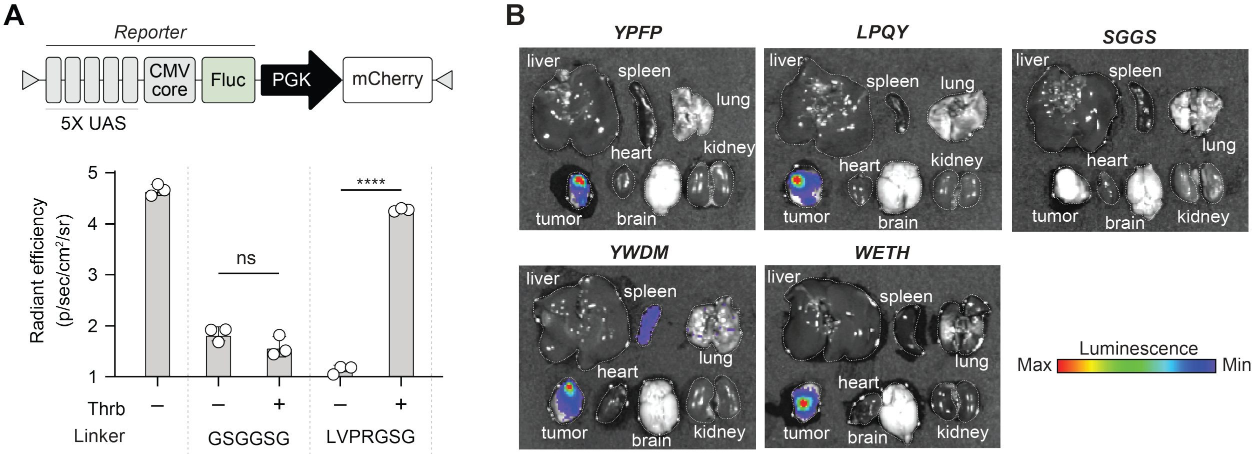


**Supplementary Figure 13. Validation of substrates identified by *in vivo* PAR T cell display.** (**A**) Gene map of PAR-activated firefly luciferase (Fluc) luminescent reporter and quantification of luminescence from primary human T cells co-expressing PAR-activated Fluc reporter and monoclonal PAR displaying either a thrombin-cleavable linker (LVPRGSG) or control linker (GSGGSG) after co-incubation with Thrb and HER2+ MDA-MB-468 cells for 24 hours at 37°C. (T-test, ****P<0.0001, $n$ = 3 biological replicates, error bars depict mean ± SD) (**B**) T cells were transduced to co-express the Fluc reporter cassette along with a monoclonal PAR encoding the indicated hit sequence (YPFP, LPQY, YWDM, or WETH) identified from *in vivo* PAR T cell display screening, or SGGS as a control linker. Twenty-four hours after injection to mice bearing HER2+ MDA-MB-468 tumors, tumor and organs were excised and submerged in luciferin as shown in representative bioluminescent images to determine biodistribution of activated PARs.

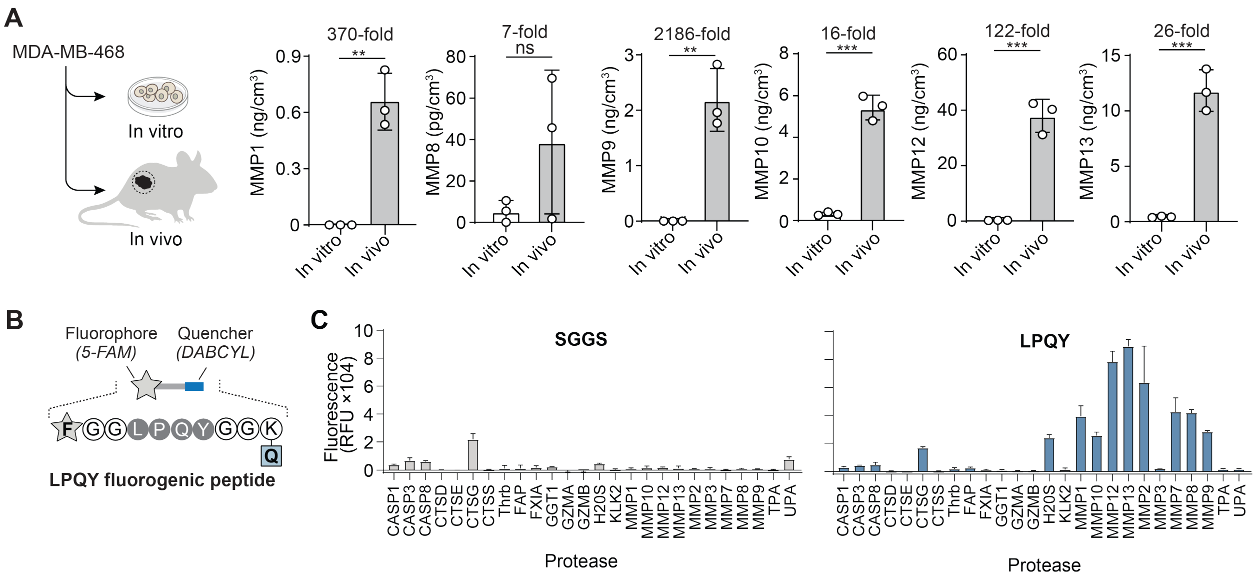


**Supplementary Figure 14. MMP expression in MDA-MB-468 HER2$^+$ cancer cells and xenograft tumors enabling LPQY substrate cleavage.** (**A**) One million cancer cells (1 mL) were cultured at 37ºC for 24 hours. Tumors were isolated and dissociated in tissue protein extraction buffer containing protease inhibitor cocktail. The supernatant was collected and analyzed by ELISA. (T-test, **P < 0.01, ***P < 0.001, *n* = 3 biological replicates, error bars depict mean ± SD). (**B**) Illustration of fluorogenic peptide design using the LPQY substrate sequence identified from *in vivo* PAR T cell display, with a fluorophore 5-FAM and quencher DABCYL pair (**C**) Bar graphs plotting the fluorescence of samples containing either the LPQY or SGGS (control) fluorogenic peptides after incubation with recombinant proteases (50 nM) for 30 min at 37°C.

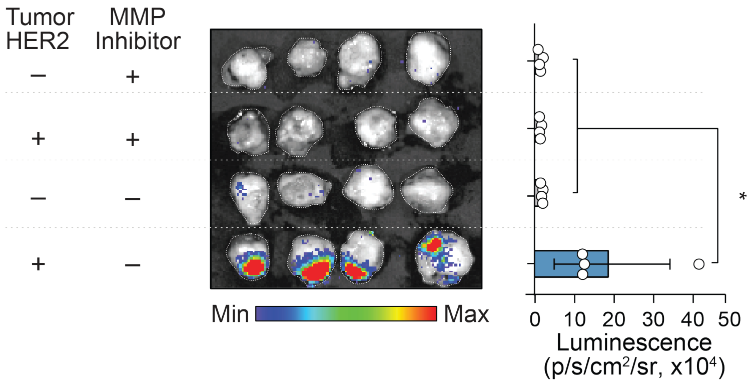


**Supplementary Figure 15. *In vivo* activation of LPQY HER2 PAR T cell sensors requires both tumor-selective proteases and antigens.** Mice bearing MDA-MB-468 HER2+ tumor were i.t. injected with the broad-spectrum matrix metalloproteinase inhibitor marimastat one day before i.v. injection of PAR T cell sensors encoding the LPQY substrate linker. Luminescence images and quantification of excised tumors were acquired 24 hours after T cell sensor injection. One-way ANOVA and Tukey post-test and correction, *$P < 0.05$, $n = 4$ biological replicates, error bars depict mean ± SD.

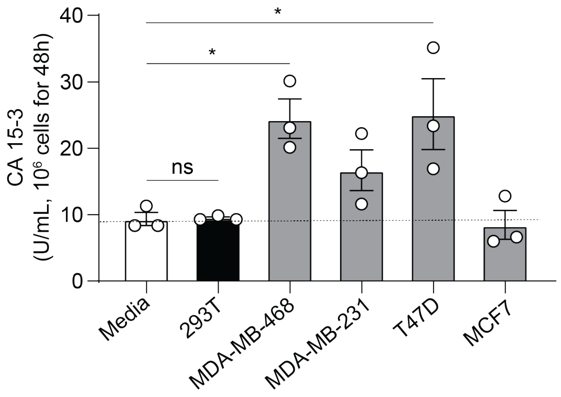


**Supplementary Figure 16. The protein biomarker CA 15-3 is secreted by human breast cancer cell lines.** Secretion level of CA 15-3 in culture media of human breast cancer cell lines including MDA-MB-468, MDA-MB-231, T47D and MCF7 and 293T (control) measured by ELISA.

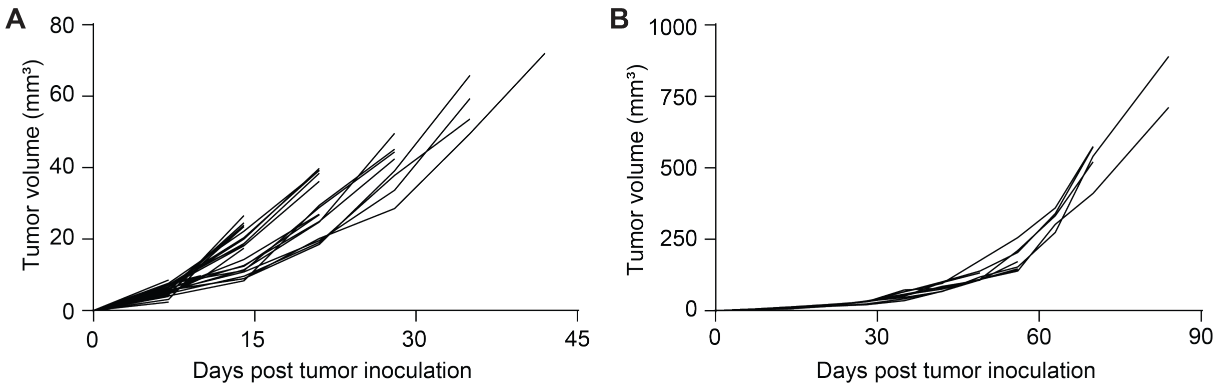


**Supplementary Figure 17. Tumor growth kinetics of MDA-MB-468 HER2+ breast cancer xenografts.** Five million MDA-MB-468 HER2$^+$ cells were subcutaneously inoculated into mice at staggered time points. Tumor volumes were measured weekly. Tumor-bearing mice (n = 38) were subsequently evaluated using OncoSCOUT to assess tumor detection performance across a range of tumor burdens, with results shown for mice bearing (**A**) <100 mm³ and (**B**) ≥100 mm³ tumors.

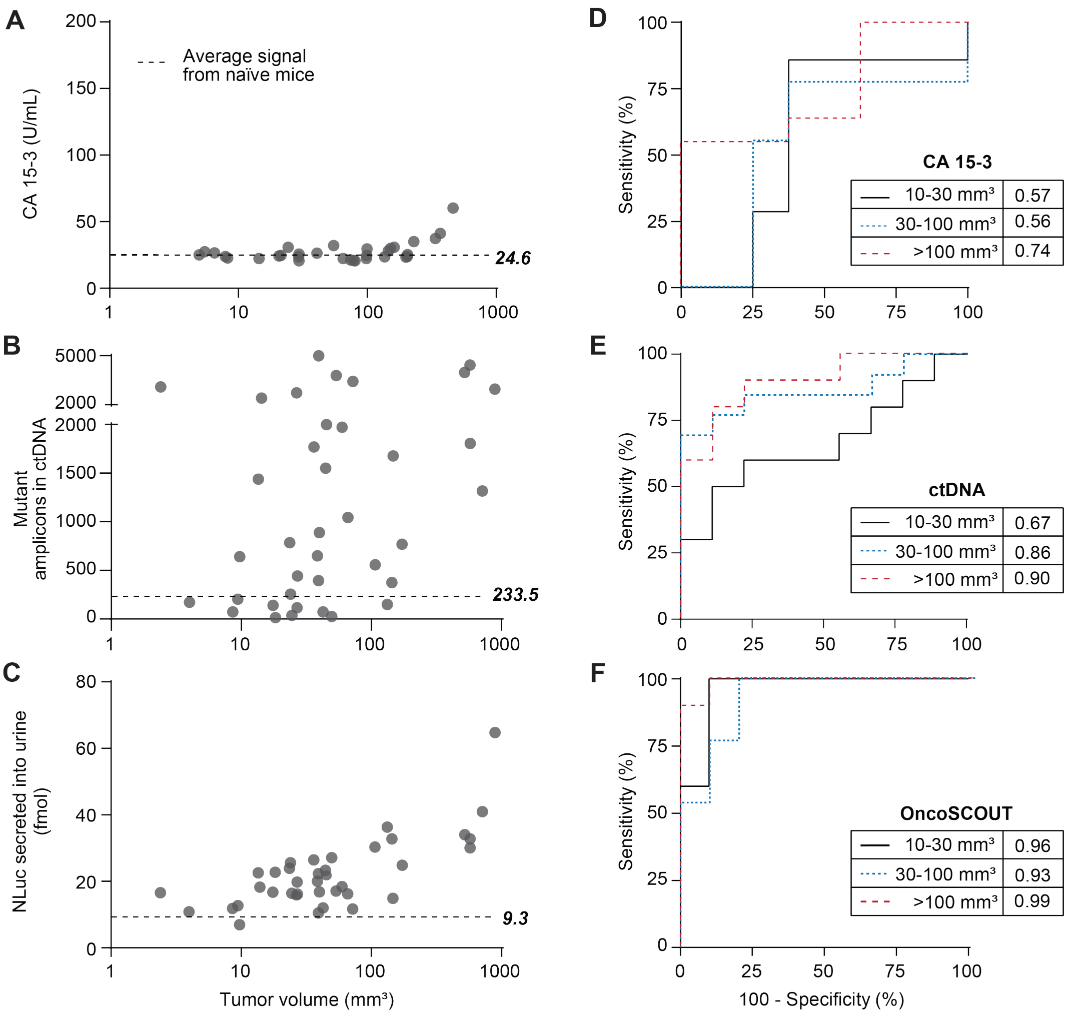


**Supplementary Figure 18. Plots of biomarker levels versus tumor size and ROC curve analyses comparing tumor detection performance of OncoSCOUT with CA 15-3 and ctDNA.** (**A**) Dot plot of serum CA 15-3 biomarker levels in naïve (dash line) and tumor-bearing mice (5-500 mm$^3$), measured by ELISA. Dashed line indicates average value for naïve mice. (**B**) Dot plot of number of mutant amplicons detected from plasma across the tumor size range (2-800 mm$^3$) same cohort with PAR T cells. Target sequencing was performed using a custom targeted gene panel. (**C**) Dot plot showing urinary NLuc signal across the tumor size range (2-800 mm$^3$). HER2 PAR T cells encoding the LPQY substrate were injected i.v. into HER2+ MDA-MB-468 tumor-bearing and naïve mice. Urine was collected, and luminescence of NLuc reporters was analyzed using an IVIS Spectrum CT system and quantified with a standard curve of recombinant NLuc. (**D**) ROC curves and AUC values using CA 15-3, (**E**) ctDNA and (**F**) OncoSCOUT, n = 5-13 biological replicates.

### Supplementary Table 1. Mutational fingerprint for MDA-MB-468 cancer cells.

| Mutation *(Gene)* | Forward Amplicon Sequence | Counts per million reads | |
|---|---|---|---|
| | | T cell | MDA-MB-468 |
| NC_000011.10: g.[108289005C>T]<br><br>*(ATM)* | *Wild-type (WT)*<br>GTATTTTTTCCCTTAACTCTGTTAGGGATTTGGATCCTGCTCCTA<br>ATCCACCTCATTTTCCATCGCATGTGATTAAAGCAACATTTGCCT<br>ATATCAGCAATTGTCATAAAACCAAGTTAAAAAGCATTTTAGAAA<br>TTCTTTCCAAAAGCCC<br><br>*Mutant (MUT)*<br>GTATTTTTTCCCTTAACTCTGTTAGGGATTTGGATCCTGCTCCTA<br>ATCCACCTTATTTTCCATCGCATGTGATTAAAGCAACATTTGCCT<br>ATATCAGCAATTGTCATAAAACCAAGTTAAAAAGCATTTTAGAAA<br>TTCTTTCCAAAAGCCC | WT: 1741<br><br>MUT: 0 | WT: 587<br><br>MUT: 588 |
| NC_000017.11: g.43092919G>A<br><br>*(BRCA1)* | *WT*<br>TGACTTTTGGACTTTGTTTCTTTAAGGACCCAGAGTGGGCAGAG<br>AATGTTGCACATTCCTCTTCTGCATTTCCTGGATTTGAAAACGGA<br>GCAAATGACTGGCGCTTTGAAACCTTGAATGTATTCTGCAGATC<br>GGAAGAGCACACGTCTGA<br><br>*MUT*<br>TGACTTTTGGACTTTGTTTCTTTAAGGACCCAGAGTGGGCAGAG<br>AATGTTGCACATTCCTCTTCTGCATTTCCTGGATTTGAAAACAGA<br>GCAAATGACTGGCGCTTTGAAACCTTGAATGTATTCTGCAGATC<br>GGAAGAGCACACGTCTGA | WT: 652<br><br>MUT: 0 | WT: 0<br><br>MUT: 281 |
| NC_000013.11: g.32355095A>G<br><br>*(BRCA2)* | *WT*<br>GGACATCCATTTTATCAAGTTTCTGCTACAAGAAATGAAAAAATG<br>AGACACTTGATTACTACAGGCAGACCAACCAAAGTCTTTGTTCC<br>ACCTTTTAAAACTAAATCACATTTTCACAGAGTTGAACAGTGTGT<br>TAGGAATATTAACTTGG<br><br>*MUT*<br>GGACATCCATTTTATCAAGTTTCTGCTACAAGAAATGAAAAAATG<br>AGACACTTGATTACTACAGGCAGACCAACCAAAGTCTTTGTTCC<br>ACCTTTTAAAACTAAATCGCATTTTCACAGAGTTGAACAGTGTGT<br>TAGGAATATTAACTTGG | WT: 1608<br><br>MUT: 0 | WT: 0<br><br>MUT: 547 |
| NC_000016.10: g.68828262C>T<br><br>*(CDH1)* | *WT*<br>TATCTTTGGCTCTCAACACTTGCTCTGTCTCCCCCACCATCCCA<br>GTTCTGATTCTGCTGCTCTTGCTGTTTCTTCGGAGGAGAGCGGT<br>GGTCAAAGAGCCCTTACTGCCCCCAGAGGATGACACCCGGGAC<br>AACGTTTATTACTATGATGA<br><br>*MUT*<br>TATCTTTGGCTCTCAACACTTGCTCTGTCTCCCCCACCATCCCA<br>GTTCTGATTCTGCTGCTCTTGCTGTTTCTTCGGAGGAGAGCGGT<br>GGTCAAAGAGCCCTTACTGCCCCCAGAGGATGACACCCGGGAC<br>AATGTTTATTACTATGATGA | WT: 2120<br><br>MUT: 0 | WT: 0<br><br>MUT: 792 |
| NC_000003.12: g.41224522A>G<br><br>*(CTNNB1)* | *WT*<br>AGTAACATTTCCAATCTACTAATGCTAATACTGTTTCGTATTTATA<br>GCTGATTTGATGGAGTTGGACATGGCCATGGAACCAGACAGAA<br>AAGCGGCTGTTAGTCACTGGCAGCAACAGTCTTACCTGGACTCT<br>GGAATCCAGATCGGAAGA<br><br>*MUT*<br>AGTAACATTTCCAATCTACTAATGCTAATACTGTTTCGTATTTGTA<br>GCTGATTTGATGGAGTTGGACATGGCCATGGAACCAGACAGAA<br>AAGCGGCTGTTAGTCACTGGCAGCAACAGTCTTACCTGGACTCT<br>GGAATCCAGATCGGAAGA | WT: 1026<br><br>MUT: 0 | WT: 0<br><br>MUT: 415 |
| NC_000001.11: g.[162780149T>C]<br><br>*(DDR2)* | *WT*<br>TCCTTTATTTTTGTTCCCAAAGACTTACCTCCCTCAACCAGCCAT<br>TTGTCCTGACTCTGTGTATAAGCTGATGCTCAGCTGCTGGAGAA<br>GAGATACGAAGAACCGTCCCTCATTCCAAGAAATCCACCTTCTG<br>AGATCGGAAGAGCACACG<br><br>*MUT*<br>TCCTTTATTTTTGTTCCCAAAGACTTACCTCCCTCAACCAGCCAT<br>TTGTCCTGACTCTGCGTATAAGCTGATGCTCAGCTGCTGGAGAA<br>GAGATACGAAGAACCGTCCCTCATTCCAAGAAATCCACCTTCTG<br>AGATCGGAAGAGCACACG | WT: 1422<br><br>MUT: 0 | WT: 534<br><br>MUT: 627 |

| Mutation *(Gene)* | Forward Amplicon Sequence | Counts per million reads | |
|---|---|---|---|
| | | **T cell** | **MDA-MB-468** |
| NC_000017.11: g.[39723662_ 39723912del]<br>*(ERBB2)* | *WT*<br>CAGATGCGGATCCTGAAAGAGACGGAGCTGAGGAAGGTGAAGG<br>TGCTTGGATCTGGCGCTTTTGGCACAGTCTACAAGGTCAGGGC<br>CAGGTCCTGGGGTGGGCGGCCCCAGAGGATGGGGGCGGTGC<br>CTGGAGGGGTGTGGTCGGCAGTTC<br>*MUT*<br>CAGATGCGGATCCTGAAAGAGACGGAGCTGAGGAAGGTGAAGG<br>TGCTTGGATCTGGCGCTTTTGGCACAGTCTACAAGGGCATCTGG<br>ATCCCTGATGGGGAGAATGTGAAAATTCCAGTGGCCATCAAAGT<br>GTTGAGGGAAAACACATCCC | WT: 794<br>MUT: 0 | WT: 334<br>MUT: 558 |
| NC_000002.12: g.[211420620_ 211420623del]<br>*(ERBB4)* | *WT*<br>CTCAGCATCCATCATATCTTCCAAATCCTCTTCATCCAAGAGATT<br>CTGAAAGAACTTGCTGTCATTTGGACTGGGAAGCTTCATACGAT<br>CATCACCCTAAAAGAAAGATTGCCCATCAGACACAAATATGAAG<br>ATCGGAAGAGCACACGTC<br>*MUT*<br>CTCAGCATCCATCATATCTTCCAAATCCTCTTCATCCAAGAGATT<br>CTGAAAGAACTTGCTGTCATTTGGACTGGGAAGCTTCATACGAT<br>CATCACCCTAAAAGATTGCCCATCAGACACAAATATGAAGATCG<br>GAAGAGCACACGTCTGAA | WT: 1880<br>MUT: 0 | WT: 372<br>MUT: 343 |
| NC_000013.11: g.28034322T>C<br>*(FLT3)* | *WT*<br>GCAGTTCTGCAGATAGAGGAAAGAATAATGAATTTTTACCTTTGC<br>TTTTACCTTTTTGTACTTGTGACAAATTAGCAGGGTTAAAACGAC<br>AATGAAGAGGAGACAAACACCAATTGTTGCATAGAATGAGATGT<br>TGTCTTGGATAGATCGG<br>*MUT*<br>GCAGTTCTGCAGATAGAGGAAAGAATAATGAATTTTTACCTTTGC<br>TTTTACCTTTTTGTACTTGTGACAAATCAGCAGGGTTAAAACGAC<br>AATGAAGAGGAGACAAACACCAATTGTTGCATAGAATGAGATGT<br>TGTCTTGGATAGATCGG | WT: 1019<br>MUT: 0 | WT: 0<br>MUT: 376 |
| NC_000002.12: g.[208248468G>A]<br>*(IDH1)* | *WT*<br>ACATTATTGCCAACATGACTTACTTGATCCCCATAAGCATGACGA<br>CCTATGATGATAGGTTTTACCCATCCACTCACAAGCCGGGGGAT<br>ATTTTTGCAGATAATGGCTTCTCTGAAGACCGTGCCACCCAGAA<br>TATTTCGTATGGTGCCAT<br>*MUT*<br>ACATTATTGCCAACATGACTTACTTGATCCCCATAAGCATGACGA<br>CCTATGATGATAGGTTTTACCCATCCACTCACAAGCCGGGGGAT<br>ATTTTTGCAGATAATGGCTTCTCTGAAGACCGTACCACCCAGAA<br>TATTTCGTATGGTGCCAT | WT: 1284<br>MUT: 0 | WT: 229<br>MUT: 286 |
| NC_000002.12: g.[208243593C>T]<br>*(IDH1)* | *WT*<br>AGGAACTGTGTGCAAAATCTTCAATTGACTTATCTTGATTATACA<br>TCCCCATGGCAACACCACCACCTTCTGTAGAGGAGAAGCCAGT<br>GAGAGGAAAAAAGGGAGAAGAATAAATGTAATGTAGTTGTAATA<br>AGGCTAAAATCACCCACCA<br>*MUT*<br>AGGAACTGTGTGCAAAATCTTCAATTGACTTATCTTGATTATACA<br>TCCCCATGGCAATACCACCACCTTCTGTAGAGGAGAAGCCAGT<br>GAGAGGAAAAAAGGGAGAAGAATAAATGTAATGTAGTTGTAATA<br>AGGCTAAAATCACCCACCA | WT: 2065<br>MUT: 0 | WT: 402<br>MUT: 442 |
| NC_000004.12: g.54736599G>C<br>*(KIT)* | *WT*<br>ACGTCTGGTCCTATGGGATTTTTCTTTGGGAGCTGTTCTCTTTAG<br>GTAAAATGATCCTTGCCAAAGACAACTTCATTAGACTCAGAGCAT<br>CTTCTTGAAGTTTCATTGGTGTCCTGCTTCCTTGTGATTAACACT<br>GCTTTGCAAACTGAGA<br>*MUT*<br>ACGTCTGGTCCTATGGGATTTTTCTTTGGGAGCTCTTCTCTTTAG<br>GTAAAATGATCCTTGCCAAAGACAACTTCATTAGACTCAGAGCAT<br>CTTCTTGAAGTTTCATTGGTGTCCTGCTTCCTTGTGATTAACACT<br>GCTTTGCAAACTGAGA | WT: 1058<br>MUT: 0 | WT: 0<br>MUT: 410 |

| Mutation *(Gene)* | Forward Amplicon Sequence | Counts per million reads | |
|---|---|---|---|
| | | **T cell** | **MDA-MB-468** |
| NC_000017.11: g.31358461G>T<br><br>*(NF1)* | *WT*<br>GGCACATTATTCTGGGGAATGTATATTATGTTTTCCACTACATAT<br>TTTCATTTAATTTTCCTCTAAAATGTTCCTCTGTTGACTTTTTTTTT<br>CTTTTAGGCATAATTTGTTGGACTCTAAGATCAACACCCTGTTAT<br>CATTGTGCCAAGAT<br><br>*MUT*<br>GGCACATTATTCTGGGGAATGTATATTATGTTTTCCACTACATAT<br>TTTCATTTAATTTTCCTCTAAAATGTTCCTCTGTTTACTTTTTTTTT<br>CTTTTAGGCATAATTTGTTGGACTCTAAGATCAACACCCTGTTAT<br>CATTGTGCCAAGAT | WT: 1402<br><br>MUT: 0 | WT: 0<br><br>MUT: 598 |
| NC_000022.11: g.29639032G>A<br><br>*(NF2)* | *WT*<br>TCAAACTGTAAAATCACAACTTTCAGGAAATGTGATAAATTTAGT<br>GGGAAAAAAATTTAATGCACGCCTTGCAAAGGCTTCTTTGAGGG<br>TAGCACAGGAGGAAGTGCCAATATAGTGTGTTTGTCTTTTGCTC<br>TGCAATTCTGCAGGTACT<br><br>*MUT*<br>TCAAACTGTAAAATCACAACTTTCAGGAAATGTGATAAATTTAGT<br>GGGAAAAAAATTTAATGCACGCCTTGCAAAGGCTTCTTTGAGGA<br>TAGCACAGGAGGAAGTGCCAATATAGTGTGTTTGTCTTTTGCTC<br>TGCAATTCTGCAGGTACT | WT: 496<br><br>MUT: 0 | WT: 0<br><br>MUT: 597 |
| NC_000009.12: g.[136503105C>G]<br><br>*(NOTCH1)* | *WT*<br>AACTCGGACGGCAACGCTCACACCCGTGGGTAGCAACTGGCAC<br>AAACAGCCAGCGTGTCTGGGGCACGGGGGGATGGCACCCCCT<br>GCAGGCAGAGCCTGTTCCCGGGATGGGGCCACACTTACTCTGC<br>ACGGCCTCGATCTTGTAGGGGAT<br><br>*MUT*<br>AACTCGGACGGCAACGCTCACACCCGTGGGTAGCAACTGGCAC<br>AAAGAGCCAGCGTGTCTGGGGCACGGGGGGATGGCACCCCCT<br>GCAGGCAGAGCCTGTTCCCGGGATGGGGCCACACTTACTCTGC<br>ACGGCCTCGATCTTGTAGGGGAT | WT: 1911<br><br>MUT: 0 | WT: 434<br><br>MUT: 376 |
| NC_000003.12: g.179204485_ 179204486delinsAA<br><br>*(PIK3CA)* | *WT*<br>TGTGCATATGTGTATGTTGAGTGTATACATTAGTATATACATACT<br>TTTTTCTTTTAGATCTATGTTCGAACAGGTATCTACCATGGAGGA<br>GAACCCTTATGTGACAATGTGAACACTCAAAGAGTACCTTAGAT<br>CGGAAGAGCACACGTCT<br><br>*MUT*<br>TGTGCATATGTGTATGTTGAGTGTATACATTAGTATATAAATACTT<br>TTTTCTTTTAGATCTATGTTCGAACAGGTATCTACCATGGAGGAG<br>AACCCTTATGTGACAATGTGAACACTCAAAGAGTACCTTAGATC<br>GGAAGAGCACACGTCT | WT: 1700<br><br>MUT: 0 | WT: 0<br><br>MUT: 610 |
| NC_000009.12: g.[95476865A>G]<br><br>*(PTCH1)* | *WT*<br>GAAAAACATCATCCACACCAACACCAAGAGCGAGAAATGGCAAA<br>ACCTACAGCAAAAACAGAGGATGGTGGCATTAGACATGCGAGAT<br>GCAATTCAGATGATTCTAAAGCTAGTTAGGACTCTGCAGATCGG<br>AAGAGCACACGTCTGAACT<br><br>*MUT*<br>GAAAAACATCATCCACACCAACACCAAGAGCGAGAAATGGCAAA<br>ACCTACAGCGAAAACAGAGGATGGTGGCATTAGACATGCGAGA<br>TGCAATTCAGATGATTCTAAAGCTAGTTAGGACTCTGCAGATCG<br>GAAGAGCACACGTCTGAACT | WT: 901<br><br>MUT: 0 | WT: 193<br><br>MUT: 222 |
| NC_000009.12: g.[95516690T>C]<br><br>*(PTCH1)* | *WT*<br>TCTTTGTCTCCCCTGTCGTCTTTTTCTTCTCCTCCGTTTTCTTCTT<br>CTTCTTCTCCTCCTCCTCCGTCTTTACAAAAGGAACGGAAAGTG<br>TAAAAACCCCGGCGCGCTGGGCCGCCGGAGGCTTTCGGCGGA<br>GTGCAGCGCGGACTCACAA<br><br>*MUT*<br>TCTTTGTCTCCCCTGTCGTCTTTTTCTTCTCCTCCGTTTTCTTCTT<br>CTTCTCCTCCTCCTCCTCCGTCTTTACAAAAGGAACGGAAAGTG<br>TAAAAACCCCGGCGCGCTGGGCCGCCGGAGGCTTTCGGCGGA<br>GTGCAGCGCGGACTCACAA | WT: 444<br><br>MUT: 0 | WT: 80<br><br>MUT: 86 |

| Mutation *(Gene)* | Forward Amplicon Sequence | Counts per million reads | |
|---|---|---|---|
| | | **T cell** | **MDA-MB-468** |
| NC_000009.12: g.[95453540G>A]<br><br>*(PTCH1)* | *WT*<br>TGATTTCTAAAACATGTCTCCTTGCACACGCCTGCTTACCTGACA ATGAAGTCGAACTCAGATCCCGCCAGCATCAGCACTCCCAGCA GAGTGGACACGGCGCCATCCAGGACGGGTGCAAACATGTGCTC CAGGGCAAGCACAGCCCTGC<br><br>*MUT*<br>TGATTTCTAAAACATGTCTCCTTGCACACGCCTGCTTACCTGACA ATGAAGTCGAACTCAGATCCCGCCAGCATCAGCACTCCCAGCA GAGTGGACACGGCACCATCCAGGACGGGTGCAAACATGTGCTC CAGGGCAAGCACAGCCCTGC | WT: 2789<br><br>MUT: 0 | WT: 589<br><br>MUT: 547 |
| NC_000010.11: g.87931090G>T<br><br>*(PTEN)* | *WT*<br>CTTCCTAAGTGCAAAAGATAACTTTATATCACTTTTAAACTTTTCT TTTAGTTGTGCTGAAAGACATTATGACACCGCCAAATTTAATTGC AGAGGTAGGTATGAATGTACTGTACTATGTTGTATAACTTAAACC CGATAGACTGTATCT<br><br>*MUT*<br>CTTCCTAAGTGCAAAAGATAACTTTATATCACTTTTAAACTTTTCT TTTAGTTGTGCTGAAAGACATTATGACACCGCCAAATTTAATTGC AGAGTTAGGTATGAATGTACTGTACTATGTTGTATAACTTAAACC CGATAGACTGTATCT | WT: 1116<br><br>MUT: 0 | WT: 0<br><br>MUT: 1145 |

Mutations use HGVS nomenclature. Brackets indicate a mutation in only one allele.

Counts per million reads were determined by sequencing the genomic DNA of primary human T cells or MDA-MB-468 cancer cells using the CleanPlex OncoZoom Cancer Hotspot Kit. Values shown are mean of two technical replicates.

WT = wild-type amplicon, MUT = mutant amplicon
